# GPR15 and CD38 define a subset of peripheral blood pathogenic effector Th2 cells associated with active eosinophilic esophagitis

**DOI:** 10.1101/2025.11.16.688265

**Authors:** Caitlin M. Burk, Duncan M. Morgan, Jonathan N. Glickman, Yamini V. Virkud, Emily Liao, Elizabeth G. Moseley, Karina Canziani, Moumita Bhowmik, Jennifer C. Li, Tarun Keswani, Navneet K. Virk-Hundal, Aubrey J. Katz, Qian Yuan, Sarita U. Patil, J. Christopher Love, Wayne G. Shreffler

## Abstract

Eosinophilic esophagitis (EoE) is a chronic allergic disease driven by exposure to culprit antigens. Due to the local nature of the inflammation, diagnosis and assessment are limited to invasive procedures. Based on prior single-cell RNA sequencing (scRNA-seq) data linking peripheral GPR15+ pathogenic effector Th2 (peTh2) cells to esophageal tissue peTh2s, we hypothesized the direct involvement of GPR15+ peTh2 cells in EoE pathogenesis and aimed to further evaluate their association with EoE disease status. We subjected samples from subjects with or without EoE to flow cytometry (n = 74 peripheral blood, 17 biopsy) and scRNA-seq (n = 27 peripheral blood, 10 biopsy). Expression of GPR15 by peripheral peTh2 cells was increased in EoE, and these cells expressed increased CD38 in active EoE--findings recapitulated in esophageal biopsies. We also identified a peTh2-associated, CD38-containing gene expression program that peripheral GPR15+ peTh2 cells upregulated in active EoE. The level of upregulation was distinct from other circulating peTh2 cells and was more similar to that seen in esophageal peTh2 cells. An association between expression of GPR15 by peripheral peTh2 cells, the aryl hydrocarbon receptor was strongest in subjects with EoE, suggesting an environmental exposure or susceptibility. The magnitude of GPR15 expression by peripheral peTh2 cells could effectively in discriminate active EoE from no EoE in our study population (AUC 0.93). Our data suggest that EoE-related peTh2 cells are identifiable and accessible in the peripheral blood, and could be exploited in both clinical practice as a non-invasive biomarker and continued investigation into mechanisms driving EoE.

**One sentence summary:** GPR15 marks a subset of peripheral blood pathogenic effector Th2 cells associated with eosinophilic esophagitis (EoE) that upregulate CD38 during active disease – an observation that has potential to be used for non-invasive diagnosis and monitoring of EoE and that has suggests new mechanisms driving this increasingly prevalent allergic disease.

## INTRODUCTION

Eosinophilic esophagitis (EoE) is a chronic, antigen-mediated allergic disease estimated to affect 1 in 700 Americans.(*1*) Suppression of eosinophil-predominant type 2 inflammation is necessary to prevent long-term consequences including fibrosis and strictures, but eosinophil-targeting medications have not met symptomatic endpoints.(*2*) This apparent gap in understanding the critical pathogenic mediator of EoE limits the development of effective and targeted treatments. Determination of EoE status is also limited by this focus on tissue-resident eosinophils--histological enumeration of eosinophils in esophageal biopsies is required for diagnosis and for monitoring responses to treatment, necessitating repeated endoscopies for patients with EoE. This requirement likely contributes to significant delays (*3*) and geographic disparities (*4*) in EoE diagnosis and imposes a significant burden of time, procedures, and cost when managing EoE.

Several studies using single-cell RNA sequencing (scRNA-seq) have identified highly polarized pathogenic effector Th2 (peTh2) cells in the esophagus in active, but not quiescent, EoE.(*5–7*) These cells are characterized by increased Th2 cytokine production, expression of receptors for IL-25, TLSP, and IL-33, and increased lipid metabolism.(*8–11*) Notably, these cells are a major source of IL-4 and IL-13, the targets of dupilumab, the only biologic medication currently approved for treatment of EoE. In human subjects, increased peTh2 cells have been described in affected tissues or multiple tissue-specific allergic diseases (*12–14*) and in the peripheral blood in eosinophilic gastroenteritis.(*9, 15*) The expansion of allergen-specific, phenotypically similar “Th2A” cells has also been described in IgE-mediated allergy.(*10, 16*)

We previously identified a rare subset of peTh2 cells in the peripheral blood of subjects with EoE that share T cell receptor (TCR) sequences with clonally-expanded esophageal peTh2 cells. These cells strongly express GPR15, a chemokine receptor previously described as involved in T cell migration to the colon and skin,(*17–22*) compared to other peripheral blood peTh2 cells.(*6*) Based on demonstrated expression of GPR15L/*C10ORF99* by the esophageal epithelium, we hypothesized that peTh2 cells may use GPR15 to migrate to the esophagus in EoE.(*6*) The exact role of GPR15 in EoE, including its expression pattern and regulation, has not been established.

Here, we investigated the GPR15+ peT2h population in the peripheral blood using flow cytometry and scRNA-seq. We found that expression of GPR15 and the activation marker CD38 by peTh2 cells can differentiate between disease states in EoE, and that expression of AhR and CD38 is related to expression of GPR15 by peripheral blood peTh2 cells. We further demonstrated that these peripheral blood GPR15+ peTh2 cells in EoE are transcriptionally and phenotypically more similar to esophageal peTh2s than to other circulating peTh2 cells. These findings lay the groundwork for a clinically translatable blood test for EoE, underlie a strategy for studying the cells that mediate EoE via the peripheral blood, and suggest a mechanism by which AhR-activating environmental exposures may influence EoE.

## RESULTS

### Expression of GPR15 by peripheral blood peTh2 cells is higher in EoE

We analyzed 74 cryopreserved peripheral blood mononuclear cell (PBMC) samples from 56 subjects undergoing clinically-indicated endoscopy (n=29 samples from active EoE, 26 from EoE in remission after an elimination diet, 11 from no EoE, additional details in Methods) and from eight subjects with known IgE-mediated food allergy without a history of EoE. Fifteen subjects with EoE provided multiple samples at different time points. There were 15 longitudinal active/dietary remission pairs. (**Tables S1-S4**). PBMC samples were thawed and subjected to flow cytometry with a 22-marker panel (**Table S5**).

We identified peTh2 cells as memory CD4+ cells (CD3+CD4+CD45RA-) that dually expressed CRTh2 and CD161, followed by analysis of GPR15 expression (**Figure 1A**). Non-peTh2 Th2 cells (“conventional Th2” or convTh2) were defined as CRTh2+CD161-.

**Figure 1.**
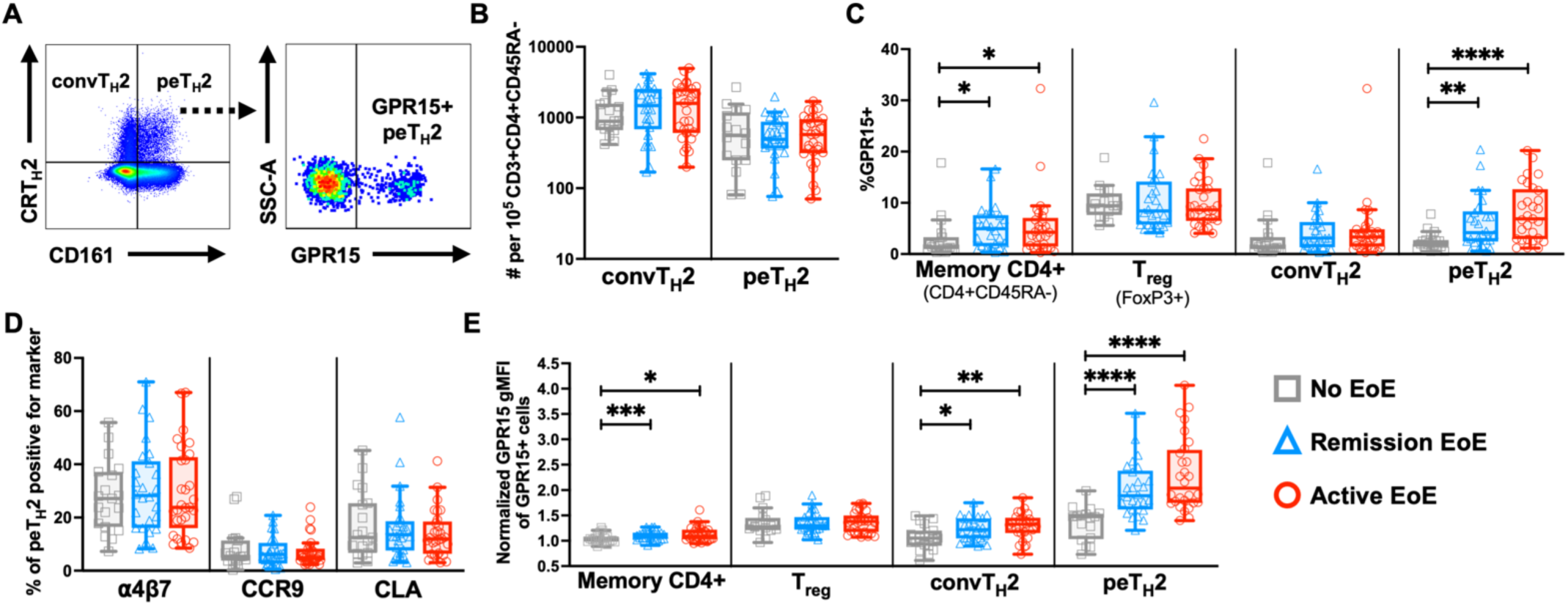
Peripheral blood peTh2 cells express more GPR15 in EoE. **(A)** Gating strategy for peripheral blood convTh2s, peTh2s, and GPR15+ peTh2s, starting from Live, CD3+CD4+CD45RA- cells. **(B)** Number of peripheral blood convTh2s and peTh2s per 10,000 memory CD4+ cells. **(C)** Percentage of peripheral blood CD4+ T cell subsets expressing GPR15. **(D)** Percentage of peripheral blood peTh2s expressing other homing markers. **(E)** Normalized gMFI of GPR15 on GPR15+ CD4+ T cell subsets. MFI normalized to CD3+CD4-CD45RA-GPR15+ cells. Differences between disease states were evaluated using Kruskal-Wallis tests followed by Wilcoxon rank-sum tests. *p<0.05, **p<0.01, ***p<0.001, ****p<0.00001. *peTh2, pathogenic effector Th2. EoE, eosinophilic esophagitis. convTh2, conventional Th2. gMFI, geometric mean fluorescence intensity. Treg, T regulatory*.

The fraction of memory CD4+ T cells that were convTh2 or peTh2 cells was similar between disease states (**Figure 1B**), differing from previous work showing an association between EoE or eosinophilic gastroenteritis generally increased peTh2 cells.(*9, 15*) We hypothesized that the GPR15-expressing subsets might vary with EoE disease state, and, indeed, the percentage of GPR15+ peTh2 cells was significantly higher in active and remission EoE compared to no EoE (**Figure 1C**). An increased GPR15+ population was also observed in overall memory CD4+ T cells in EoE, but not in T regulatory (Treg) cells or convTh2 cells (**Figure 1B**). To determine if this pattern in peTh2 cells was unique to GPR15, we also analyzed the expression of three additional homing markers: α4β7 (gut), CCR9 (duodenum), and CLA (skin). There were no differences in expression of these markers by peTh2 cells between EoE disease states (**Figure 1D**). Most GPR15+ cells expressed one of these other homing markers, but the levels did not significantly vary between disease states. (**Figure S1A**) Within the subset of 15 paired samples, there was no difference between active and remission EoE, suggesting that GPR15+ populations may expand when EoE develops and remain a consistent size, regardless of disease status. (**Figure S1B**).

We next focused on the magnitude of GPR15 expression by GPR15+ cells, hypothesizing that, although active and quiescent EoE have a similar frequency of circulating GPR15+ peTh2 cells, elevated expression of GPR15 by GPR15+ peTh2 cells in active EoE might promote migration of GPR15+ peTh2 cells to the esophagus.

The geometric mean fluorescence intensity of GPR15 was higher in active EoE and EoE in dietary remission compared to no EoE in peTh2 cells, convTh2 cells, and memory CD4+ T cells (**Figure 1E**). In the subset of paired samples, there was also no trend in GPR15 MFI between active disease and remission (**Figure S1C**). The only difference between disease states in α4β7, CCR9, and CLA MFIs in any of the memory CD4+ subsets analyzed was a higher CLA MFI in Treg cells in not EoE compared to active EoE (**Figure S1D**). These findings may suggest an EoE-associated exposure or process leading to widespread increased GPR15 expression by CD4+ T cells that disproportionately affects peTh2 cells.

### Active EoE is associated with expression of CD38 by GPR15+ peTh2 cells in the peripheral blood

We next examined CD38 on peTh2 cells, based on literature demonstrating increased expression of CD38 by antigen-specific Th2A cells during pollen season or after a positive oral food challenge,(*10*) suggesting that CD38 may function as an activation marker for peTh2-like populations. In other contexts, expression of CD38 is increased in esophageal—but not peripheral blood—CD4+ T cells in active EoE,(*23*) and in peripheral blood, gut-homing, tetramer-positive effector T cells in celiac disease after gluten exposure.(*24, 25*)

We found no difference in the proportion of convTh2 cells or peTh2 cells expressing CD38 between disease states. (**Figure 2A**) The percentage of GPR15+ peTh2 cells expressing CD38 was, however, significantly higher in active EoE compared to EoE in dietary remission and no EoE (**Figures 2B, D**). Expression of CD38 by GPR15+ peTh2 cells was also significantly higher in active disease compared to remission in the subset of paired samples (**Figure 2C**).

**Figure 2.**
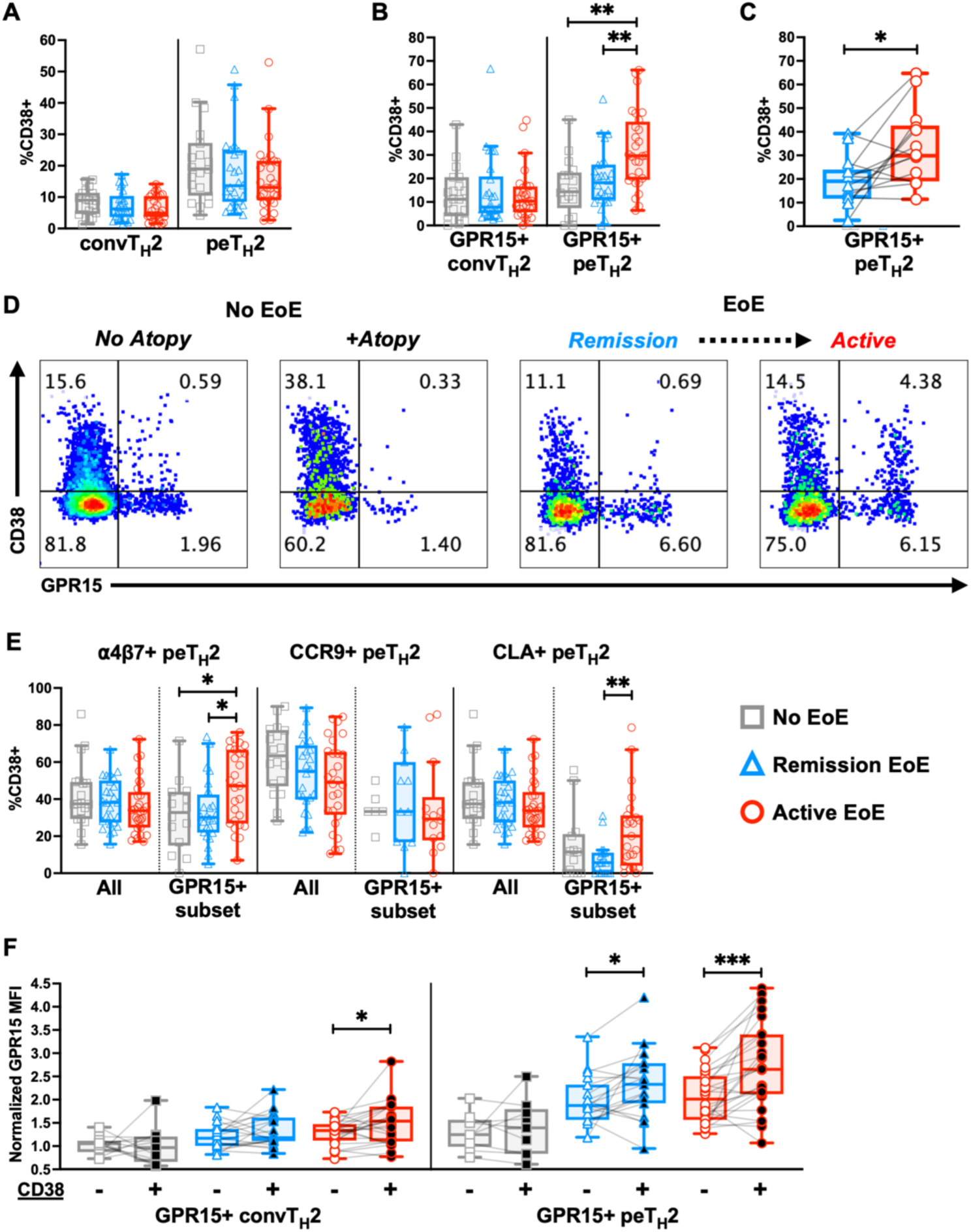
A greater proportion of peripheral blood GPR15+ peTh2 cells express CD38 in active EoE. Percentage of **(A)** overall peripheral blood convTH2s and peTH2s, and **(B)** peripheral blood GPR15+ convTH2s and GPR15+ peTH2s expressing CD38. **(C)** Percentage of peripheral blood GPR15+ peTH2s expressing CD38 in longitudinal samples. **(D)** Representative plots of GPR15 and CD38 expression by peripheral blood peTH2s in individuals with and without EoE. The EoE samples are from the same subject over time. The non-EoE subject with atopy had a history of asthma, allergic rhinitis, eczema, and IgE-mediated food allergy. **(E)** CD38 positivity in peripheral blood peTH2s expressing ⍺4β7, CCR9, and CLA, with or without GPR15. **(F)** Normalized gMFI of GPR15 on GPR15+ convTH2s and GPR15+ peTH2s that were CD38+ or CD38-. MFI is normalized to CD3+CD4-CD45RA-GPR15+ cells. Lines connect cell populations in the same samples. Population differences between disease states were evaluated using a Kruskal-Wallis test followed by Wilcoxon rank-sum tests. Paired samples were evaluated with Wilcoxon signed-rank tests. **\***p<0.05, **p<0.01, ***p<0.001. *peTH2, pathogenic effector TH2. EoE, eosinophilic esophagitis. convTH2, conventional TH2. gMFI, geometric mean fluorescence intensity*.

Expression of CD38 was not different between disease states in the subsets of peTh2 cells expressing α4β7, CCR9, and CLA (**Figure 2E**). In the subsets of GPR15+ peTh2 cells co-expressing the other examined homing markers, only α4β7+GPR15+ peTh2 cells had a similar pattern to GPR15+ peTh2 cells (**Figure 2E**).

Within GPR15+ peTh2 cells in EoE, cells expressing CD38 had a higher GPR15 MFI compared to cells that were CD38- (**Figure 2F**). This relationship was not present in samples from subjects without EoE. GPR15 MFI was also higher in CD38+GPR15+ convTh2 cells compared to CD38-GPR15+ convTh2 cells in active disease (**Figure 2F**). MFI of other gut-homing markers on peTh2 cells increased with CD38 positivity, but this phenomenon was not always specific to EoE, suggesting that CD38+ peTh2 cells may be poised to better migrate to the gut in general (**Figure S2**).

### convTh2 and peTh2 gene expression programs are present among peTh2 cells in the peripheral blood in EoE

To assess transcriptional features associated with GPR15+ peTh2 cells in the peripheral blood, we analyzed scRNA-seq data generated from flow-sorted peTh2, convTh2, and CD4+ memory non-Th2 cells (CD45RA-CRTh2-) isolated from PBMCs following *ex vivo* stimulation for 6 hours with anti-CD3/anti-CD28-coated beads (**Figure 3A**). We compiled transcriptional data from 27 samples collected from 21 subjects (n = 14 active EoE, 13 EoE in dietary remission; characteristics of previously unpublished samples in **Table S6**; data from eight of these subjects were previously reported.(*6*)

**Figure 3.**
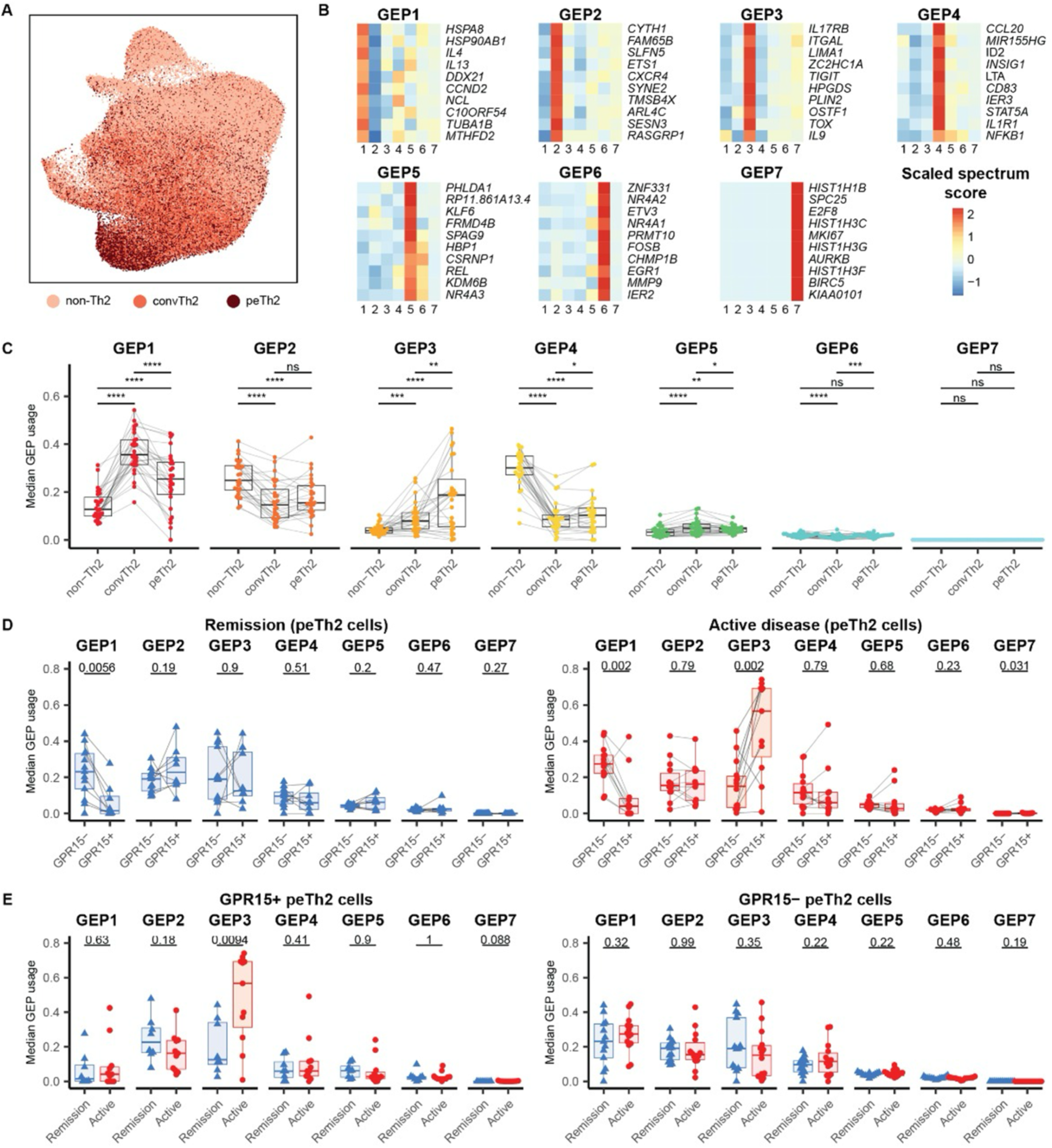
Single-cell analysis of peripheral blood CD4+ T cells in EoE reveals upregulation of a peTh2-associated GEP in GPR15+ peTh2s in active disease. **(A)** UMAP of ex vivo stimulated non-Th2, convTh2, and peTh2 from peripheral blood. **(B)** Scaled gene spectrum scores for the top ten genes in each GEP inferred using consensus non-negative matrix factorization. **(C)** Median normalized usage of each GEP among non-Th2, convTh2, and peTh2 cells from each patient. **(D)** Median normalized usage of each GEP among GPR15+ and GPR15- peTh2 cells from samples with active disease and remission. **(E)** Median normalized usage of each GEP among GPR15+ and GPR15- peTh2 cells samples with active disease and remission. All samples shown have greater than one GPR15+ convTh2 cell. P-values are calculated with a two-sided t-test and are adjusted with Bonferroni correction. ****p<0.0001. *EoE, eosinophilic esophagitis. GEP, gene expression program. peTH2, pathogenic effector TH2, UMAP, uniform manifold approximation projection. convTH2, conventional TH2*.

To define functionalities present among *ex vivo* stimulated peTh2 cells, we applied an approach for gene expression program (GEP) inference based on consensus non-negative matrix factorization.(*26, 27*) This analysis identified seven distinct, reproducible programs present among peTh2 cells (**Figure 3B**, **Figure S3A, Supplementary data file**). To assess whether these programs were specific to peTh2 cells, we determined the usage of each of these programs among convTh2 and non-Th2 cells and compared these values with those of peTh2 cells.(*28*) We found that two programs, GEP1 and GEP3, exhibited substantially increased usage by CRTh2+ cell subsets. Specifically, usage of GEP1 was greatest among convTh2 cells, and usage of GEP3 was greatest among peTh2 cells (**Figure 3C**). Consistent with these trends, the genes contributing most robustly to GEP1 included transcripts associated with a convTh2 phenotype, including the cytokines *IL4* and *IL13*, while transcripts with the greatest contributions to GEP3 included features associated with a peTh2 phenotype, including *IL17RB*, *HPGDS*, and *IL9.* (**Figure 3B**, **Supplementary data file**) (*8–10*) Of GEP1 and GEP3, CD38 only exhibited substantial contribution to GEP3, suggesting a strong relationship between these pathogenic features of peTh2 cells and the expression of CD38. (**Figure S3B, Supplementary data file**) GEP3 also included genes related to lipid metabolism and eicosanoid synthesis that have been implicated in EoE (*PPARG, PLA2G4A, PTGS2, ALOX5AP*) and several cytoskeletal proteins, similar to the signature we previously identified in esophageal-resident peTh2s in active EoE.(*6*)

Two other GEPs, including GEP2, which included transcripts broadly associated with T cells (*PTPRC*, *CXCR4*) and GEP4, which contained TH1- (*IFNG*, *TNF*, *IL2*) and TH17- (*CCL20*, *IL17A*, *RORC*) associated transcripts, were expressed in convTh2 and peTh2 cells albeit at levels lower than in non-Th2 cells, (**Figure 3C**) suggesting that they represent programs that may be used by Th2 cells but are not specific to this subset. The three remaining GEPs exhibited usage confined to smaller subsets of cells. GEP5 included features associated with NF-κB signaling (*REL*, *NFKBIA*), GEP6 included features associated with early T cell activation (*NR4A2*, *FOS*, *JUNB*), and GEP7 was used by very few cells and included features associated with cycling cells (*HIST1H1B*, *HIST1H3C*, *MK167*), in addition to CD38. (**Figure 3C**, **Supplementary data file**) In total, this analysis identifies a collection of gene expression programs utilized by peTh2 cells, including a program for convTh2 features (GEP1) and an additional program for peTh2 features (GEP3).

### GPR15+ peTh2 cells upregulate a peTh2-specific gene program compared to other peripheral blood peTh2 cells

We next examined the usage of these GEPs in peTh2 cells with or without detectable expression of GPR15. The frequency of GPR15 expression in this single-cell data was lower than that determined by flow cytometry, (**Figure S3C**) likely due to incomplete recovery of lowly-expressed transcripts.(*29*) Strikingly, we found that peTh2 cells with detected GPR15 transcript from patients with active disease, but not remission, exhibited significantly upregulated use of GEP3, the program most strongly associated with peTh2 features. (**Figure 3D**) This finding was also manifest in conventional Th2 cells in active disease, (**Figure S4A**) perhaps suggesting, along with the observed decrease in GEP1 usage in active but not remission EoE, that these cells may be more activated or becoming more polarized. When peTh2 cells were stratified by other homing markers, including *ITGA4*, *SELPLG*, or *ITGA7*, they did not display differences between positive and negative groups.(**Figure S4B-E**) We also compared the usage of each GEP between disease statuses and found that GPR15+ peTh2 cells—but not GPR15- peTh2 cells—exhibited increased usage of GEP3 in active disease compared to remission. (**Figure 3E**) Similar trends were not observed with convTh2s. (**Figure S4F**) Together, these results suggest that GPR15 is a marker of a subset of peTh2 cells in peripheral blood that exhibit increased polarization toward a peTh2 phenotype in subjects with EoE.

### GPR15 expression by peripheral blood peTh2 cells in EoE is associated with the aryl hydrocarbon receptor

We next examined transcription factors known to promote expression of GPR15: AhR, GATA3, FoxP3, and RORγt. (*18, 20, 30*) In the scRNA-seq dataset, peripheral blood peTh2 cells expressed GATA3 and AhR, but negligible FoxP3 or RORγt. (**Figure S5A**) Expression of transcription factors was similar between active EoE and dietary remission. (**Figure S5A**) The transcript for *AHRR*, an AhR regulator and AhR-responsive target gene, (*31*) was upregulated in GPR15+ peTh2 cells compared to GPR15- peTh2 cells in active EoE, but not in remission. (**Figure 4A**) *AHRR* was also upregulated in active disease versus dietary remission in GPR15+ peTh2 cells, but not in GPR15- peTh2 cells, suggesting that these cells may exhibit increased AhR activity. (**Figure 4B**) Notably, in the GEP inference analysis, *AHRR* contributed the most to GEP3, the GEP associated with GPR15+ peTh2 cells in active EoE. (**Figure S3B**)

**Figure 4.**
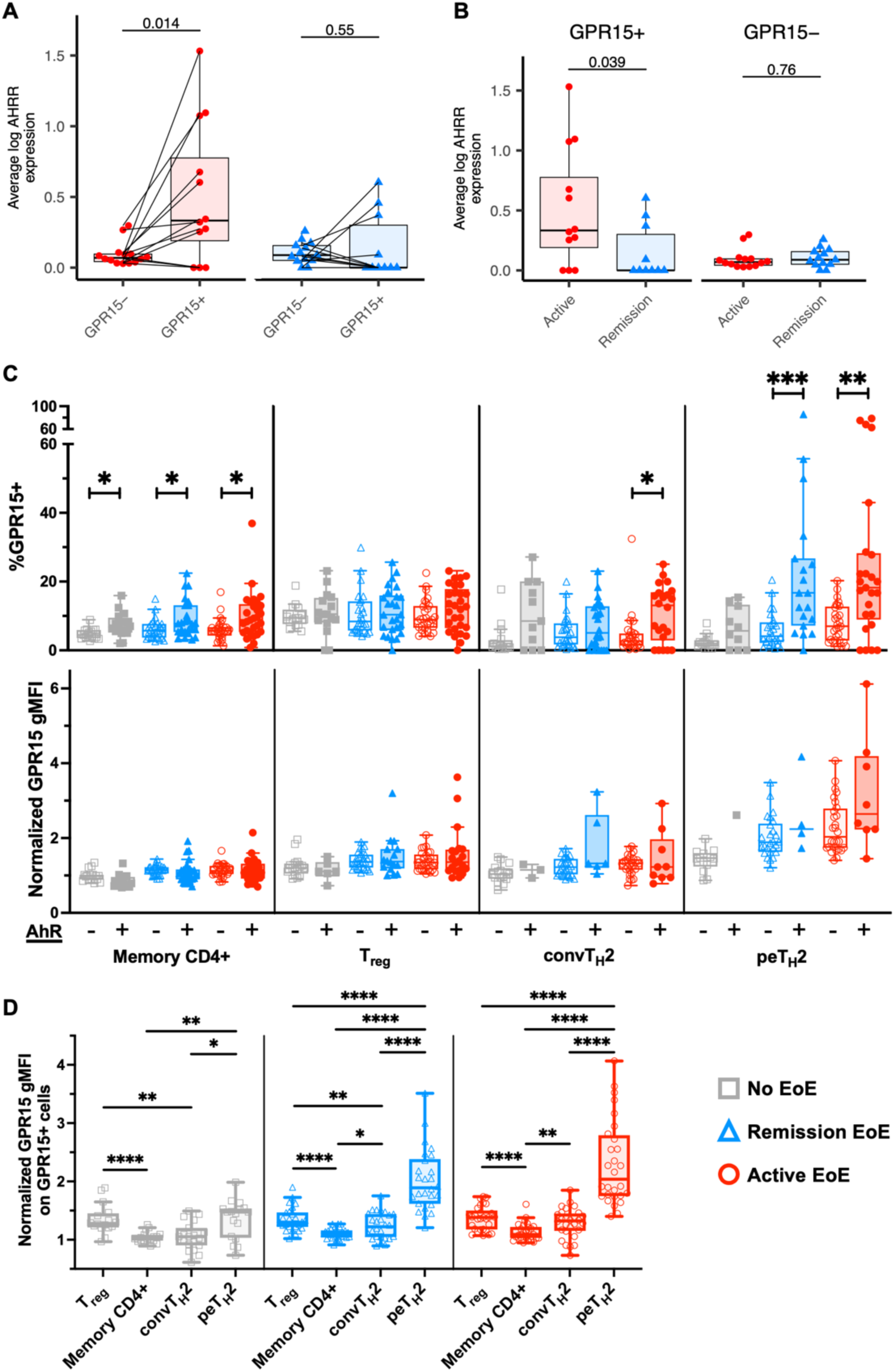
GPR15 expression by peripheral blood peTh2 cells in active EoE is associated with AhR. **(A, B)** Average pseudobulk expression of *AHRR* among GPR15+ and GPR15- peTh2 cells. P-values are calculated with a two-sided t-test. **(C)** Percentage of cells expressing GPR15 (top) and normalized GPR15 gMFI (bottom) of AhR+ and AhR- CD4+ memory T cell subsets. **(D)** Normalized GPR15 gMFI in Tregs, the overall memory CD4+ population, convTh2 cells, and peTh2 cells. MFI is normalized to CD3+CD4-CD45RA-GPR15+ cells. Wilcoxon rank-sum tests were used to evaluate differences between populations in **(C)** and **(D)**. *p<0.05, **p<0.01, ***p<0.001, ****p<0.0001. *peTh2, pathogenic effector Th2. EoE, eosinophilic esophagitis. convTH2, conventional Th2. gMFI, geometric mean fluorescence intensity*.

By flow cytometry, AhR+ cells were more likely to be GPR15+ in peTh2 cells in both active and remission EoE, in convTh2 cells in active EoE only, and in all disease states in memory CD4+ cells (**Figure 4C**). GPR15 MFI on GPR15+ cells was not significantly associated with AhR expression in any cell types, (**Figure 4C**) but there was a trend towards increased GPR15 MFI in AhR+ peTh2s. (The very small number of peTh2 cells co-expressing AhR and GPR15 in our dataset limits our analysis here, however.)

We next reexamined GPR15 MFI, comparing memory CD4+ subsets within each EoE disease state to understand patterns of GPR15 upregulation that may be common to all subjects or specific to EoE. In all conditions, the overall memory CD4+ subset had the lowest GPR15 MFI. In unaffected subjects, convTh2 cells had a similar MFI to memory CD4+ cells, and FoxP3+ and peTh2 cells had the highest, and similar, MFIs. (**Figure 4D**) In subjects with EoE in remission after an elimination diet, GPR15 MFI on convTh2 and peTh2 cells was higher, with convTh2 cells surpassing memory CD4+ cells and peTh2 cells surpassing FoxP3+ cells. (**Figure 4D**) Finally, in active EoE, the GPR15 MFI on convTh2 cells was no longer significantly lower than that on FoxP3+ cells. (**Figure 4D**) This pattern may suggest that FoxP3+ and peTh2 cells are predisposed to express increased amounts of GPR15 at baseline via different regulatory mechanisms, and that the potential for peTh2 and convTh2 cells to express GPR15 is enhanced in EoE.(*8*)

### Circulating GPR15+ peTh2 cells are transcriptionally similar to esophagus-resident peTh2 cells

We next determined the usage of each of the transcriptional GEPs in T cells present in the esophagus during EoE in our previously-published scRNA-seq data (n = 6 with active EoE, 4 with EoE in dietary remission).(*6*) As expected, we observed increased usage of GEP1 (convTh2 features) and GEP3 (peTh2 features) in esophagus-resident peTh2 cells relative to other esophageal T cell subsets, (**Figure S6A**) confirming that these programs are also associated with peTh2 cells that likely migrate to the esophagus in active EoE.

We then compared the usage of each GEP by GPR15- peTh2 cells and GPR15+ peTh2 cells in the peripheral blood with peTh2 cells detected in the esophagus during active EoE. Usage of GEP1 was elevated and usage of GEP3 was reduced among GPR15- peTh2 cells relative to either of the other subsets. Strikingly, we detected no significant differences in the usage of any of the GEPs between GPR15+ cells in the peripheral blood and peTh2 cells detected in the esophagus (**Figure 5A**), indicating a substantial degree of phenotypic similarity between these two populations. This finding implies that peTh2 cells present in the esophagus in EoE are highly similar to GPR15+ peTh2 cells in the peripheral blood and could indicate that GPR15+ peTh2 peripheral blood cells are an intermediate population on a trajectory towards the most polarized pathogenic esophageal peTh2 phenotype in EoE.

**Figure 5.**
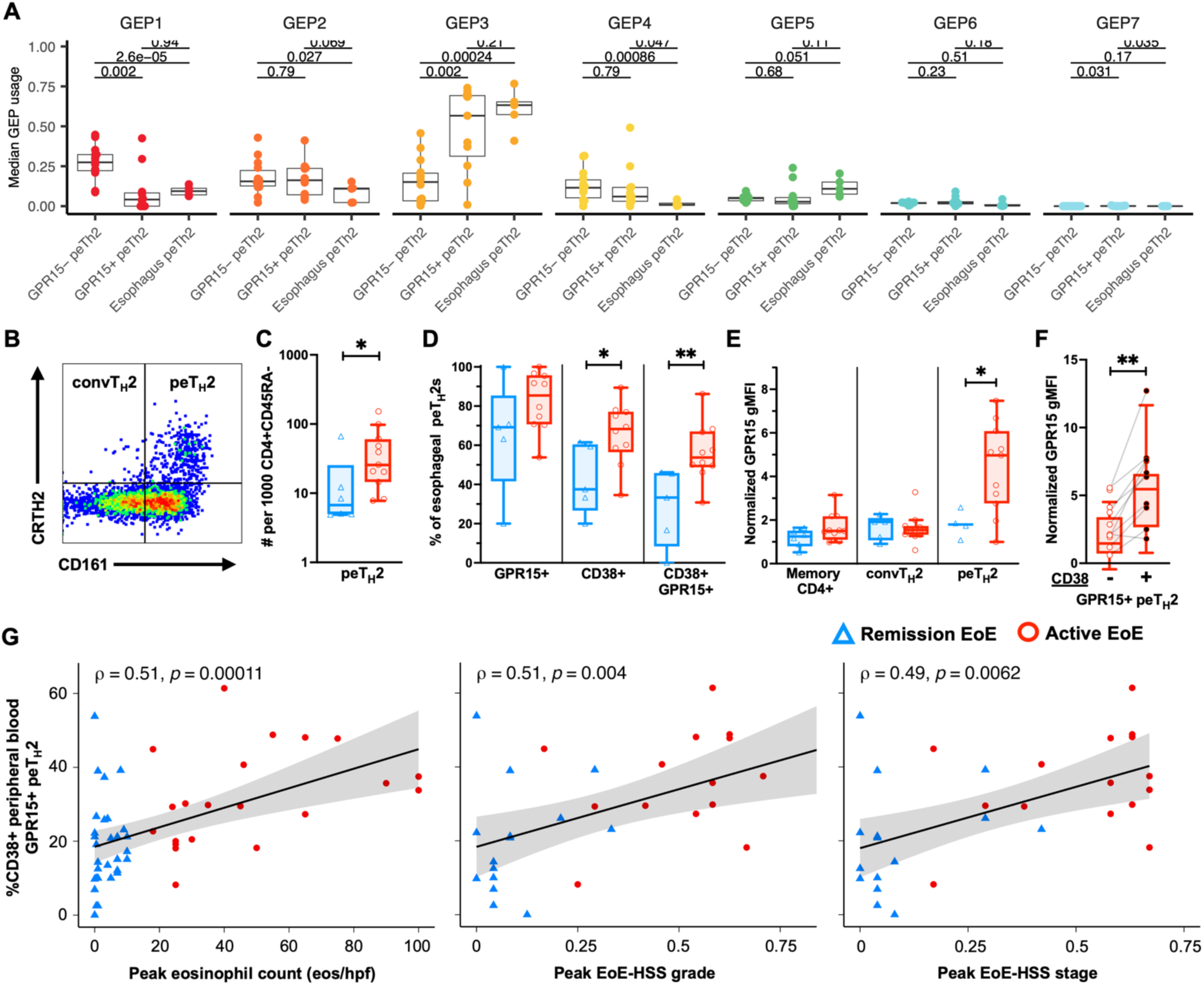
Circulating GPR15+CD38+ peTh2 cells in EoE are highly similar to esophageal peTh2 cells and correlate with histology. (**A**) Median normalized usage of each GEP among circulating GPR15- and GPR15+ peTh2 cells, and esophageal peTh2 cells from each patient. P-values are calculated with a two-sided t-test and are adjusted with Bonferroni correction. **(B)** Gating strategy for esophageal convTH2 cells and peTH2 cells, starting from Live, CD3+CD4+CD45RA- cells. (**C**) Number of esophageal peTh2 cells per 1000 memory CD4+ T cells. (**D**) Percentage of esophageal peTh2 cells expressing GPR15, CD38, or both markers. **(E)** Normalized gMFI of GPR15 on esophageal T cell subsets. MFIs are normalized to CD3+CD4-CD45RA-GPR15+ cells. (**F**) Normalized gMFI of GPR15 on GPR15+ peTH2 cells that were CD38+ or CD38-. Lines connect cell populations in the same samples. For flow cytometry data, population differences between disease states were evaluated using Kruskal-Wallis tests followed by Wilcoxon rank-sum tests. Paired samples were evaluated with Wilcoxon signed-rank tests. (**G**) Spearman correlations and 95% confidence interval for the percentage of peripheral blood GPR15+ peTh2 cells that were CD38+ and esophageal histology measures in EoE samples. *p<0.05, **p<0.01. *peTH2, pathogenic effector TH2. EoE, eosinophilic esophagitis. GEP, gene expression program. convTH2, conventional TH2. gMFI, geometric mean fluorescence intensity. EoE-HSS, EoE histologic scoring system*.

### Expression of GPR15 and CD38 by esophageal peTh2 cells in EoE mirrors patterns observed in peripheral blood peTh2 cells

We next assessed phenotypic similarity between peripheral blood and esophageal peTh2 cells by cytometry. Cells from fresh esophageal biopsies from 17 subjects undergoing clinically-indicated endoscopies (n = 11 with active EoE, 6 with EoE in remission after an elimination diet, **Table S7**) were disassociated and analyzed by flow cytometry (see **Methods, Table S8**) for expression of GPR15 and CD38 by peTh2 cells. (**Figure 5B**)

The proportion of CD4+ T cells that were peTh2 cells was higher in active EoE, but peTh2 cells were present in samples from remission.(**Figure 5C**) In active EoE, a higher proportion of peTh2 cells were CD38+ and GPR15+CD38+, and there was a trend towards an increase in GPR15+ cells (**Figure 5C**)—a pattern similar to what we observed in peTh2 cells in the peripheral blood and consistent with previous findings regarding CD38 expression in esophageal biopsies in EoE.(*23*) Distinct from our findings in the peripheral blood, expression of GPR15 by esophageal peTh2 cells was higher in active EoE compared to remission, (**Figure 5E**) suggesting that high expression of GPR15 may only aid in migration of peTh2 cells to the esophagus during active EoE, or that expression of GPR15 by esophageal peTh2 cells is only maintained in the setting of active EoE. Esophageal peTh2 cells expressing CD38 did, similar to the circulating population, also express increased levels of GPR15 in active EoE. (**Figure 5F**) (Assessment of the relationship between CD38 and GPR15 in samples from subjects in remission was limited by cell number.) These findings support the notion that the GPR15+CD38+ peTh2 subset in the peripheral blood is closely related to the major disease-associated esophageal peTh2 population in EoE.

### GPR15 and CD38 expression by peripheral blood peTh2s correlate with histological measures of EoE disease state

Given the relationship between peripheral blood GPR15+ peTh2 cells and esophageal peTh2 cells in EoE, we next investigated the relationship between GPR15 and CD38 expression by peripheral blood peTh2 cells and EoE-related histological findings in the esophagus. A subset of 59 of the peripheral blood samples had paired esophageal biopsies with quantified peak eosinophil counts, and 42 sets of paired, same-day clinical biopsies had EoE histologic scoring system (EoE-HSS) scores.(*32*) Spearman correlation was used to compare a subset of peripheral blood measures (percent of peTh2 cells that were GPR15+, percent of GPR15+ peTh2 cells that were CD38+, GPR15 MFI on GPR15+ peTh2 cells) with esophageal eosinophil count and EoE-HSS scores. (**Figure S7**)

In the subset of subjects with known EoE, the percentage of GPR15+ peTh2s that were CD38+ positively correlated with nearly all histological components of EoE severity (**Figure 5G, S7**). There were moderate correlations among GPR15+CD38+ peTh2 cells and peak esophageal eosinophil count and EoE-HSS scores (**Figure 5G**). All analyzable EoE HSS components were also either weakly or moderately correlated with expression of CD38 by GPR15+ peTh2 cells (**Figure S7**). When subjects were further subdivided into active EoE or EoE in dietary remission, other factors were more correlated with histological measures. In active EoE but not remission, GPR15 MFI correlated with esophageal eosinophil count. (**Figure S7**) In EoE in remission but not active disease, the percentage of peTh2s that were GPR15+ correlated with all three histological measures. (**Figure S7**) This correlation of expression of GPR15 and CD38 on peripheral blood peTh2 cells with the majority of EoE-associated esophageal histological features reinforces the likelihood that peripheral blood GPR15+ peTh2 cells are key players in the pathogenesis of EoE.

### Expression of GPR15 and CD38 by peripheral blood peTh2 cells can discriminate between EoE disease states

Building on the associations we identified between EoE status and expression of GPR15 and CD38 by peripheral blood peTh2 cells, we next evaluated the discriminatory power of our observations in the peripheral blood by generating receiver-operator characteristic (ROC) curves for all peripheral blood measures associated with active disease versus no EoE or versus EoE in remission after an elimination diet (**Figure 6A**, **Table S9, Supplementary data file**). (Given the possible interdependency of GPR15 and CD38 expression, multiple-factor ROC curves were not generated.) Area under the curve (AUC) values ranged from 0.69-0.93. (**Table S9)** An example of a potential translatable use of peripheral blood peTh2 cell GPR15 MFI (AUC = 0.93) is shown in **Figure 6B**. In our cohort, 21 of 45 samples fell either above the 100% sensitivity or below the 100% specificity cut points. If these cut points were validated in an independent, large cohort, approximately half of patients with suspected EoE might be spared the need for endoscopic biopsies. The diagnostic AUC in our cohort approaches or outperforms initial diagnostic AUCs reported for proteins detected via the esophageal string test (0.97 for MBP1 and CLC/Gal-10, <0.9 for EDN, EPX, ECP),(*33*) a less-invasive diagnostic currently in clinical use for monitoring of EoE.

**Figure 6.**
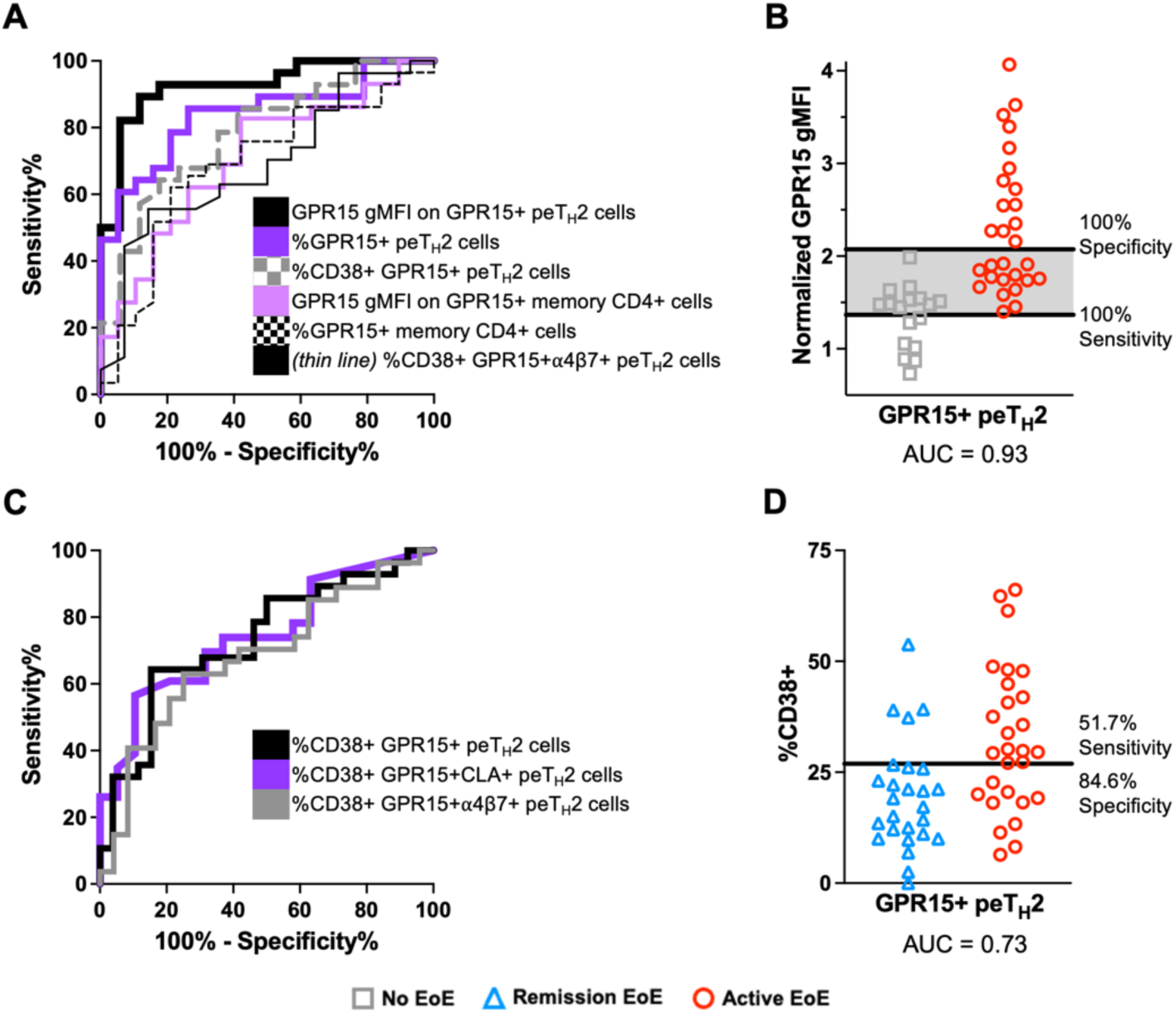
Diagnostic potential of flow cytometry for peripheral blood GPR15+ peTh2 cells in EoE. (**A**) ROC curves for factors discriminating samples from subjects with active EoE from subjects without EoE. (**B**) GPR15 gMFI on GPR15+ peTh2 cells. MFIs are normalized to CD3+CD4-CD45RA-GPR15+ cells. (**C**) ROC curves for factors discriminating samples from subjects with active EoE from subjects with EoE in remission after dietary elimination. Paired samples were considered independent in the ROC analysis. (**D**) Percentage of GPR15+ peTh2 cells that are CD38+, with line indicating the cut point discussed in the text. *peTH2, pathogenic effector TH2. EoE, eosinophilic esophagitis. ROC, receiver-operator characteristic. gMFI, geometric mean fluorescence intensity. AUC, area under the curve*.

This approach can also be applied in distinguishing between active disease and remission after an elimination diet in known EoE (**Figure 6C**). Area under the curve (AUC) values ranged from 0.68-0.74. (**Table S9)** A possible cut point (28.3%, sensitivity = 57.1%, specificity = 84.6%) for the percentage of GPR15+ peTh2 cells that were CD38+ is shown in **Figure 6D**. Patients with values above this cut point might be considered “likely active” and could proceed to additional food elimination without an endoscopy. Patients below the cut point could be considered indeterminate and undergo endoscopy to evaluate disease status. Using this strategy in our cohort, a treatment decision might be considered without an endoscopy in 22 out of 54 (41%) subjects —18 of 28 (64%) subjects with active EoE might be spared an endoscopy, but 4 of 26 (15%) subjects in remission might proceed to possibly unnecessary treatment escalation.

## DISCUSSION

We evaluated an EoE-associated subset of peripheral blood peTh2 cells in subjects with active EoE, EoE in remission induced by an elimination diet, and without EoE. This study builds on our previous findings that peripheral blood peTh2 cells with TCR sequences matching those found in esophageal peTh2 cells express the chemokine receptor GPR15 in EoE.(*6*)

We initially identified that, in subjects with EoE, an increased percentage of peripheral blood peTh2 cells express an increased amount of GPR15 compared to subjects without EoE. Given GPR15’s role in T cell migration in other inflammatory diseases, (*21, 22, 34, 35*) the expression of GPR15L by esophageal epithelium in EoE,(*6*) and the relative lack of esophageal peTh2s in remission, (*5–7*) we had hypothesized that expression of GPR15 on circulating GPR15+ peTh2 cells would be higher during active EoE compared to remission. The similar GPR15 levels we found in both EoE disease may suggest that the GPR15+ peTh2 population expands when EoE is established and remains poised to react if an antigen is reintroduced, and spurred us to investigate other aspects of peTh2s that might differ between active disease and remission.

We subsequently showed that CD38, an activation marker expressed by circulating antigen-specific Th2A cells in IgE-mediated allergy after antigen exposure,(*10*) was increased on circulating and esophageal GPR15+ peTh2 cells in active EoE. In our scRNA-seq data, CD38 was also associated with the peTh2-related GEP3, which contains additional integrins (*ITGAL*) that may synergize with GPR15 to facilitate migration when upregulated. Whether CD38 also actively facilitates migration via its known capacity to interact with CD31 (PECAM-1) on endothelium, (*36, 37*) or whether it solely marks cells endowed with the ability to migrate is not discernible with our observational data. Combining this observation with our data relating CD38 expression to higher GPR15 on peTh2 cells, we hypothesize that when EoE is established or transitions from remission to active, TCR stimulation by antigen induces both GPR15 and CD38 in a subset of allergen-specific peTh2 cells, which expand and migrate to the esophagus. In remission due to an elimination diet, the stimulus inducing CD38 is withdrawn and GPR15+ peTh2 cells are no longer able to efficiently migrate, but remain available to respond to antigen when reintroduced.

Perhaps most significantly from a clinical perspective, the level of expression of GPR15 by circulating GPR15+ peTh2 cells appears to discriminate effectively between active EoE and no EoE with excellent test performance in our cohort. Translating this potential 6-7 color flow cytometry test (CD3, CD4, CD45RA, CRTh2, CD161, GPR15, viability marker if necessary) to patients requires validation in a larger population, but it may represent an effective, non-invasive, and accessible method for diagnostic screening for EoE without an endoscopy. As noted above, the performance of this biomarker for diagnosis is comparable to that of the esophageal string test, which currently requires an office visit and is approved for EoE monitoring after diagnosis only. Previous work has also found that a combination of several eosinophil-related factors in the plasma can distinguish between untreated EoE and non-EoE controls,(*38*) but the method (relying on multiple ELISAs) has not yet been commercialized. Compared to other T-cell-based approaches in development,(*39*) our potential method does not require manipulation or stimulation of cells beyond staining for flow cytometry.

Differentiating between active and remission EoE with GPR15+CD38+ peTh2 cells on a population basis was less effective, but our data in longitudinal samples suggest that it may be possible to track the status of individual patients over time. Additional work examining the population of GPR15+CD38+ cells over time through various disease states and different time points after starting an intervention is likely to derive additional measures that might be used to determine disease status.

Our work also highlights a method to identify peripheral blood T cells associated with a tissue-specific, food-driven disease without prior knowledge of the trigger food. Though most “remission” subjects in this study responded to milk elimination, there were a total of nine different food triggers in our cohort of patients in remission, and many of the subjects in active disease have not yet identified their triggers. With our group’s previously published work(*6*) and the findings presented here, we have shown the feasibility of identifying disease-associated peripheral T cells with disease-related surface markers identified by scRNA-seq and TCR clonotype analysis of paired peripheral blood and affected tissue samples rather than using activation-induced marker assays (*40, 41*) or MHCII:peptide tetramers.(*42*) Studying peripheral blood GPR15+ peTh2s in EoE may serve as a generalizable model for studying other allergic diseases through disease-specific surface markers expressed by peTh2s initially identified through unsupervised sequencing methods.

The antigen-agnostic nature of our approach may, in fact, allow us to more readily identify triggers of EoE in the future. Currently, the only options for determining food triggers are empiric elimination diets followed by endoscopic evaluation or esophageal string test to determine if EoE is in remission, or a milk-specific 6-day PBMC stimulation assay on a research basis.(*43*) With the identification of circulating GPR15+CD38+ peTh2 cells associated with active EoE, we could now take advantage of the easily-accessible nature of these cells and determine both food triggers and food-associated TCRs through further *ex vivo* evaluation.

Open mechanistic questions include the origin of peripheral blood GPR15+CD38+ peTh2 cells. Regarding their physical origin, possibilities include emigration from lymph nodes after exposure to antigen versus recirculation of activated tissue-resident esophageal peTh2 cells. Though the esophagus does not classically have tertiary lymphoid structures, the reported penetration of food into esophageal tissue in EoE, (*44, 45*) expression of MHCII by esophageal epithelial cells,(*46*) and described presence of esophageal Th2-multipotent progenitor cells (Th2-MPP) in EoE (though lower than other examined tissues (*14*)) could support the second scenario. We, however, favor an extra-esophageal origin of these peripheral blood peTh2 cells given the persistence of the peripheral blood population in remission despite very few esophageal cells, and anecdotal reports of EoE in patients receiving 100% of their nutrition via gastrostomy tube. Further study of paired esophageal and peripheral blood samples, including in those without EoE, could shed light on this question.

In terms of the ontologic origin of peripheral GPR15+ peTh2 cells, our data suggest that expression of GPR15 by peripheral blood CD4+ T cells is associated with EoE, AhR expression, and with Th2 polarization. In peTh2 cells, increased GPR15 expression related to these factors is more dramatic, and is additionally associated with CD38 expression. Expression of FoxP3 is also associated with GPR15, but in our data FoxP3+ cells did not exhibit EoE- or AhR-associated increases in GPR15, suggesting two distinct pathways leading to expression of GPR15 in memory CD4+ T cells. The increase in GPR15+ cells and GPR15 MFI in several CD4+ cell types in EoE may suggest an exposure or process specific to EoE or that can trigger EoE in susceptible subjects, and that disproportionately affects peTh2 cells. An environmental exposure would be consistent with the rapidly increasing incidence of EoE,(*47*) and AhR is poised to mediate the effect of an exposure with its ability to recognize both exogenous toxins and endogenous, microbiome-derived compounds.(*48*) These data could be consistent with previous work showing upregulation of GPR15 on CD4+ T cells in response to Ahr ligands,(*30, 49, 50*) the additive effects of GATA3 and AhR on GPR15 expression,(*18, 30*) and increased GATA3 expression by peTh2 cells compared to convTh2 cells.(*30*) Previous evidence indicating that convTh2 cells can increase GATA3 expression and differentiate into peTh2 cells with repeated TCR stimulation (*8*) may suggest that GPR15 expression could increase as cells become progressively Th2 polarized in the context of repeated antigen exposure.

We hypothesize that activation of convTh2 cells or peTh2 cells in susceptible individuals and in the presence of AhR ligands triggers engagement of both GATA3 and AhR, leading to activated, CD38+ and GPR15^hi^ peTh2 cells, which then migrate to the esophagus and mediate EoE-related inflammation (**Figure 7**). Further work is needed to elucidate the viability of targeting these processes to treat the underlying processes underlying EoE-related inflammation.

**Figure 7.**
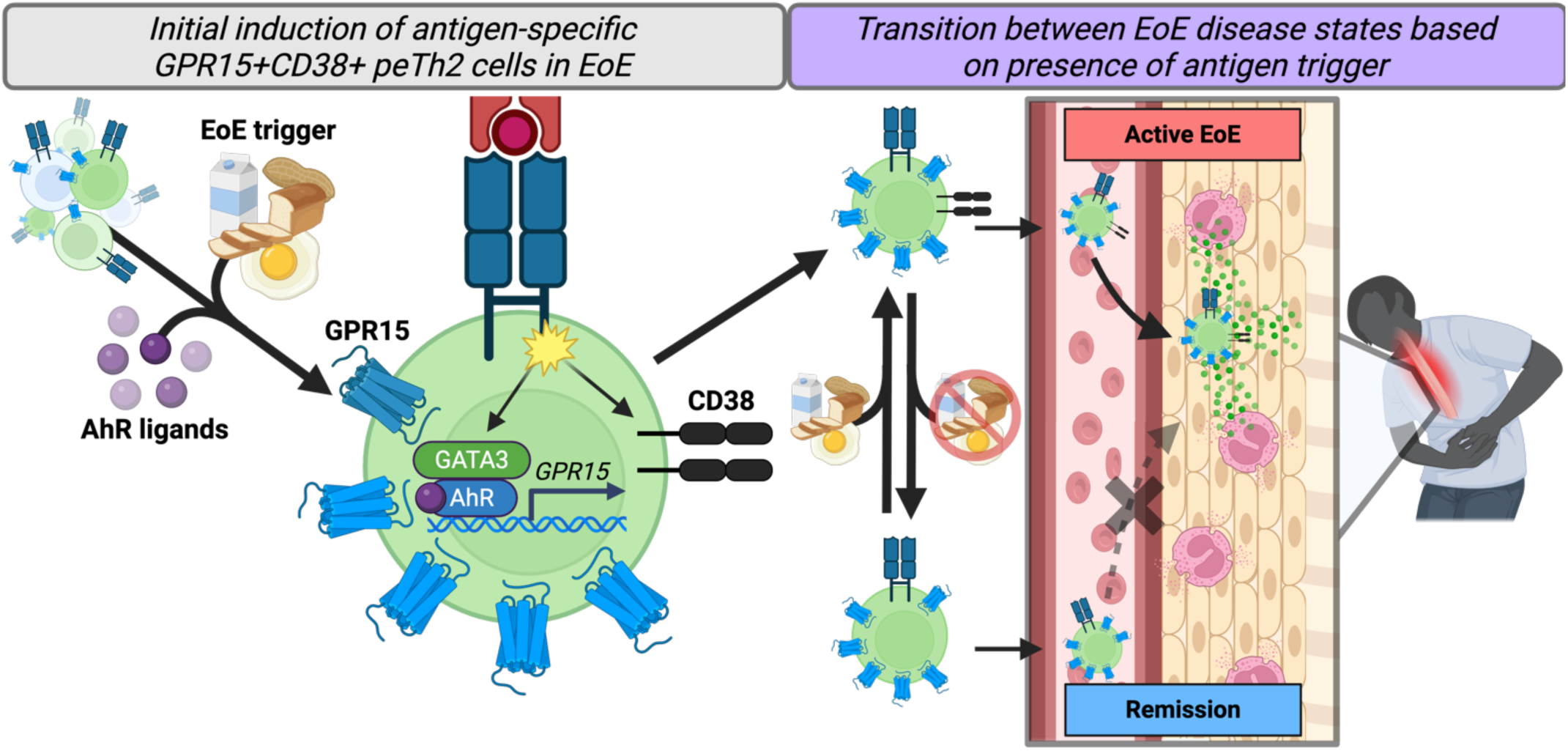
Hypothesized role of GPR15+CD38+ peTh2 cells in EoE. We hypothesize that GPR15+CD38+ peTh2 cells expressing high levels of GPR15 are generated when allergen-specific Th2 cells are exposed to their cognate antigen and AhR ligands. We posit that this leads to the expression of CD38 via TCR activation, and to high expression of GPR15 via the activation of GATA3 by TCR activation and AhR via AhR ligands. These GPR15+CD38+ peTh2 cells are then able to migrate to the esophagus and mediate EoE-related inflammation via secretion of Th2 cytokines. When the antigen is withdrawn, GPR15+ peTh2 cells decrease CD38 expression and no longer migrate to the esophagus. Reintroduction of antigen causes activation and CD38 expression. *peTh2, pathogenic effector Th2. EoE, eosinophilic esophagitis. AhR, aryl hydrocarbon receptor*.

Strengths of this study include the relatively large number of subjects included and correspondence of the majority of our samples with same-day clinical endoscopy results to verify clinical status. Limitations include our reliance on samples from patients rather than subjects enrolled in a structured clinical trial, wherein we do not have consistent lengths of time on elimination diets in our remission group (range ∼2 months to 1+ years). To enrich for food allergy in the no EoE group, we also included samples from food allergy studies that relied on clinical history to determine EoE status rather than a definitively negative biopsy. The predominance of male participants and a trend towards older participants in our EoE groups reflects the epidemiology of EoE, and likely the delay in diagnosis for many subjects, respectively. Our study population does, however, skew heavily towards white participants, which does not reflect the epidemiology of EoE (*51*) and is more likely related to disparities in access to pediatric subspecialty care, including endoscopy.

In summary, we used flow cytometry and scRNA-seq to evaluate the peripheral blood GPR15+ peTh2 population in EoE. We found that this population increases its size and expression of GPR15 in EoE, and that these cells express CD38 in active disease. These measures appear to have potential as translatable non-invasive biomarkers for EoE diagnosis and monitoring. We identified an association between AhR expression and GPR15, the pattern of which suggested a role for an EoE-related environmental exposure or sensitivity that most dramatically affects peTh2 cells. We also found that the transcriptional phenotype of peripheral blood GPR15+ peTh2 cells was more similar to esophageal peTh2 cells than to other peripheral peTh2 cells, and that expression of CD38 and GPR15 by peripheral blood peTh2 cells correlated with esophageal histological measures of disease. Overall, our results indicate that GPR15 can be used to identify peripheral blood peTh2 cells that are involved in EoE tissue-specific pathogenesis. Our findings may underlie a shift in how we can study certain immunologic processes in EoE – going forward, more accessible peripheral blood samples rather than esophageal biopsy samples might be used for evaluation of disease-modifying strategies targeting the antigen-specific or T cell mechanisms driving EoE.

## MATERIALS AND METHODS

### Sample collection from subjects undergoing endoscopy

Blood was collected immediately before upper endoscopy, during which clinical and research biopsies were obtained. PBMCs were isolated per standard Ficoll methods and cryopreserved. Biopsies were consumed within hours of collection as described below. Active EoE was defined as ≥15 eosinophils per high-power field (eos/hpf) in clinical biopsies. EoE in remission after an elimination diet was defined as <15 eos/hpf while on an elimination diet as the only treatment. Subjects on a proton pump inhibitor (PPI) were included if they had a history of an endoscopy revealing active EoE while on a PPI. No EoE was defined as <15 eos/hpf in a subject without a history of EoE and not on any treatment known to be effective in EoE (i.e., elimination diet, PPI, dupilumab, swallowed topical steroids, systemic immunosuppressants). These definitions apply to all sections of this report. Demographic information was collected via questionnaire and chart review. This study was approved by the Mass General Brigham Institutional Review Board (MGB IRB protocol nos. 2010P002087 and 2011P001159).

### Sample collection from subjects enrolled in food allergy studies

Confirmed food-allergic subjects enrolled in a food allergy biobank collecting blood samples alongside clinical blood draws or in a clinical trial evaluating intestinal permeability during two double-blind, placebo-controlled food challenges. (MGB IRB protocol nos. 2021P003078 and 2022P000392) For the former, samples from subjects who had skin prick testing or specific IgE above the 95% positive predictive value for at least one allergen were used. For the latter, samples after placebo challenges from patients who reacted during an active challenge were used. PBMCs were isolated per standard Ficoll methods and cryopreserved in liquid nitrogen. Demographic information was collected via questionnaire and chart review.

### Flow cytometry analysis of peripheral blood T cells

Cryopreserved PBMCs were thawed and then labeled with the 22-antibody panel in **Table S4**. Surface staining was performed at 4°C for 30 minutes. After fixation and permeabilization (eBioscience™ Foxp3/Transcription Factor Fixation/Permeabilization kit, Invitrogen), intracellular staining was performed at room temperature for 30 minutes. Data were acquired with a 5-laser Aurora spectral flow cytometer (Cytek) and analyzed with FlowJo v10.10 (BD Biosciences). Gating strategy and representative staining are shown in **Figures S8-9**. For percentages or gMFI determinations, only samples with five or more cells in a subset were included. N’s for each figure are shown in **Table S10**.

### Single-cell sequencing of peripheral blood T cells

Memory CD4+ cells were isolated from thawed cryopreserved PBMC samples (EasySep Human Memory CD4+ T cell Enrichment Kit, Stemcell Technologies) and cultured for 6 hours in AIM-V media (Gibco) at a concentration of 5 x 10^6^ cells per milliliter with human T-Activator CD3/CD28 beads (Thermo Fisher) in a 1:3 ratio of beads to T cells. After harvesting, cells were stained at 4°C for 30 minutes with the antibody panel in **Table S11** and/or Totalseq-A hashing antibodies (Biolegend). CD3+CD4+CD45RA-CRTH2+CD161+, CRTH2+CD161-, and CRTH2−CD161- T cells were sorted with a FACSAria II (BD Biosciences) instrument and were processed for scRNA-seq using the Seq-Well platform as previously described.(*6, 52, 53*) The Nextera XT kit (Illumina, San Diego, CA) was used for library barcoding and amplification. A Novaseq 6000 (Illumina) was used for sequencing. Hash tag oligo (HTO) libraries were amplified as previously described (Stoeckius Genome Biology) and sequenced on a Nextseq 550.

### Processing of scRNA-sequencing reads

Raw sequencing reads were demultiplexed and analyzed using the Drop-seq pipeline.(*54*) Briefly, reads were aligned to the hg38 genome and collapsed by cell barcode and unique molecular identifier. Cells with fewer than 500 unique genes were filtered out.

### Processing of cell hashing data

Cell hashing reads were aligned to HTO barcodes using CITE-seq-Count v1.4.2. (https://github.com/Hoohm/CITE-seq-Count) Cell hashing data was processed as described in Zagorulya et al.(*55*) Briefly, we first log-normalized the HTO matrix and initially partitioned the data with k-medoids clustering, with k determined by the number of multiplexed HTO barcodes within a sample. Each of these clusters corresponded to a single HTO identity. To select a positivity threshold for each HTO, we iteratively applied Grubbs’ test to the lowest value of the corresponding HTO in each cluster to remove all outliers with a threshold of p=0.05. Based on these thresholds, we then classified cells as positive or negative for each marker, and removed cells that were negative for all HTOs or positive for more than one HTO (doublets).

### scRNA-seq data processing and visualization

Previously published scRNA-seq data were obtained from GSE175930. Cells with fewer than 500 unique genes were filtered out. Data was normalized, and 2,000 variable genes were selected using the FindVariableFeatures() function in Seurat, and the ScaleData function was used to regress out the number of genes in each cell.(*56, 57*) Data was then batch corrected with Harmony to regress out patient and batch effect.(*58*) For visualization, batch corrected data underwent further dimension reduction with principal components analysis, and UMAP was performed with the top 30 principal components.

### Inference of gene expression programs with cNMF

Consensus nonnegative matrix factorization was used to identify gene expression programs present among peTh2 cells.(*26, 27*) First, batch correction was performed by applying Harmony to the single-cell count matrix (starTRAC).(*28*) Then, cNMF was performed with k-values ranging from 4 to 15; seven GEPs were selected because this number of programs maximized the stability of the resulting programs. The starCAT method was then used to predict the usage of these programs in other T cells (convTh2 cells, non-Th2 cells, peTh2 cells in the esophagus).(*26*) In analysis of GEP scores, the total program usage in each cell was normalized to one.

### Flow cytometry analysis of esophageal biopsies

Fresh esophageal biopsies were minced and digested as previously described (*6*), and individual samples were stained for 30 minutes at 4°C for 30 minutes with the antibody panel in **Table S8**. Samples were analyzed with a FACSAria III (BD Biosciences) instrument and analyzed with FlowJo v10.10.

### EoE-HSS scoring

Where available, stored clinical pathology biopsies were reevaluated for EoE-HSS scores. Scores were assigned by a single pathologist per the method described in Collins et al.(*32*)

### Statistical analysis

Statistical analysis was performed in R software v4.4.2 and GraphPad Prism v10.4.1. All boxplots are Tukey-style. Specific statistical tests are indicated in the figure legends. P-values <0.05 were considered statistically significant.

## List of supplementary materials

Figures S1-S9

Tables S1-S11

## Acknowledgements

This study would not have been possible without the patients and families who agreed to contribute samples without any foreseeable individual benefits. We are incredibly grateful for their participation. We would also like to acknowledge several generations of Shreffler laboratory technicians, including M. Kirpas, K. Gregory, J. DePaz, and A. Jaynes, for excellent technical support in processing and preserving samples. We would also like to thank the Mass General for Children Pediatric Endoscopy Center team for their assistance and teamwork in collecting samples for this study. We thank N. Kamelamela, C. Hallee, S. Levine, and the MIT BioMicro Center for assistance with library preparation and sequencing, as well as the MIT Koch Institute Flow Cytometry Core. CMB would like to thank B. Armstrong, T. Cool, and M. Dirnt for essential moral support during the writing process.

## Funding

National Institutes of Health grant T32-HL116275 (CMB, JCLi)

Consortium of Eosinophilic Gastrointestinal Disease Researchers (CEGIR) Scholar Training Award (CMB). CEGIR (U54 AI117804) is part of the Rare Disease Clinical Research Network (RDCRN), an initiative of the Office of Rare Diseases Research (ORDR), NCATS, and is funded through collaboration between NIAID, NIDDK, NCATS. CEGIR is also supported by patient advocacy groups including American Partnership for Eosinophilic Disorders (APFED), Campaign Urging Research for Eosinophilic Diseases (CURED), and Eosinophilic Family Coalition (EFC). As a member of the RDCRN, CEGIR is also supported by its Data Management and Coordinating Center (DMCC) (U2CTR002818). Funding support for the DMCC is provided by the National Center for Advancing Translational Sciences (NCATS) and the National Institute of Neurological Disorders and Stroke (NINDS).

Food Allergy Science Initiative (JCLove, WGS)

This work was supported in part by the Koch Institute Support (core) NIH Grant P30-CA14051 from the National Cancer Institute, as well as the Koch Institute-Dana-Farber/Harvard Cancer Center Bridge Project.

## Author contributions

Conceptualization: CMB, DMM, JCLove, WGS

Data Curation: CMB, DMM, EL, EGM

Formal analysis: CMB, DMM

Funding acquisition: CMB, JCLove, WGS

Investigation: CMB, DMM, JNG, CZ, MB, JCLi

Methodology: CMB, DMM, YVV, JCLove, WGS

Project administration: CMB, DMM

Resources: YVV, TK, NKV-H, AJK, QY, SUP, JCLove, WGS

Software: CMB, DMM, YVV

Supervision: JCLove, WGS

Visualization: CMB, DMM, EGM

Writing – original draft: CMB, DMM, WGS

Writing – review & editing: CMB, DMM, JNG, YVV, EL, EGM, KC, MB, JCLi, TK, NKV-H, AJK, QY, SUP, JCLove, WGS

## Competing interests

CMB, DMM, JCLove, and WGS are co-inventors on patent number PCT/US2025/016902 (METHODS AND MATERIALS TO DISTINGUISH BETWEEN ACTIVE EOSINOPHILIC ESOPHAGITIS (EoE), EoE IN REMISSION, AND NON-EoE STATES, publication date August 28, 2025) based on a portion of the findings in this manuscript.

Additional interests as follows:

CMB has received research grants from the National Institutes of Health (NIH), the Consortium of Eosinophilic Gastrointestinal Disease Researchers (CEGIR), the Campaign Urging Research for Eosinophilic Disease (CURED), and Food Allergy Research and Education, Inc (FARE). As of December 2025, she will be employed by Sanofi, but performed the work herein, including drafting and submitting the initial manuscript while employed at MGH.

JCLi is currently employed by Pfizer Inc., but performed the work herein during her fellowship training at MGH.

YVV has received research grants from the NIH.

NKV-H has received clinical trial funding from Regeneron and AstraZeneca.

QY has received research funding and/or clinical trial funding from EvoEndo, Inc, the Food Allergy Science Initiative (FASI), the Gerber Foundation, Infinant Health. He is a medical advisor for Tiny Health.

SUP has received research grants from the NIH, Food Allergy Science Initiative, and Buhlmann Laboratories. She has received clinical trial support from Regeneron and FARE. She has received consulting fees from Seismic Therapeutics and Mabylon, and royalties from Uptodate.

JCLove has interests in Sunflower Therapeutics PBC, Pfizer, Honeycomb Biotechnologies, OneCyte Biotechnologies, SQZ Biotechnologies, Alloy Therapeutics, QuantumCyte, Amgen, and Repligen.

JCLove’s interests are reviewed and managed under the Massachusetts Institute of Technology (MIT)’s policies for potential conflicts of interest. JCLove receives sponsored research support at MIT from Amgen, the Bill & Melinda Gates Foundation, Biogen, Pfizer, Roche, Takeda, and Sanofi.

WGS has received clinical trial funding from Aimmune Therapeutics, ALK, Celgene, DBV Technologies, Genentech, Novartis, Regeneron Pharmaceuticals, and Vedanta; served on scientific advisory boards for Aimmune Therapeutics and FARE; received personal consulting fees from Aimmune Therapeutics/Nestle, Harmony, Merck, and Novartis; received royalties from UpToDate; and received grants from the NIH, Demarest Lloyd Foundation, Thornhill Foundation, and FASI.

DMM, JNG, EL, EGM, KC, MB, TK, AJK declare that they have no competing interests.

## Data sharing

Data will be available in ImmPort, GEO, and dbGAP.

