## Supplemental figures for "GPR15 and CD38 define a subset of peripheral blood pathogenic effector Th2 cells associated with active eosinophilic esophagitis"

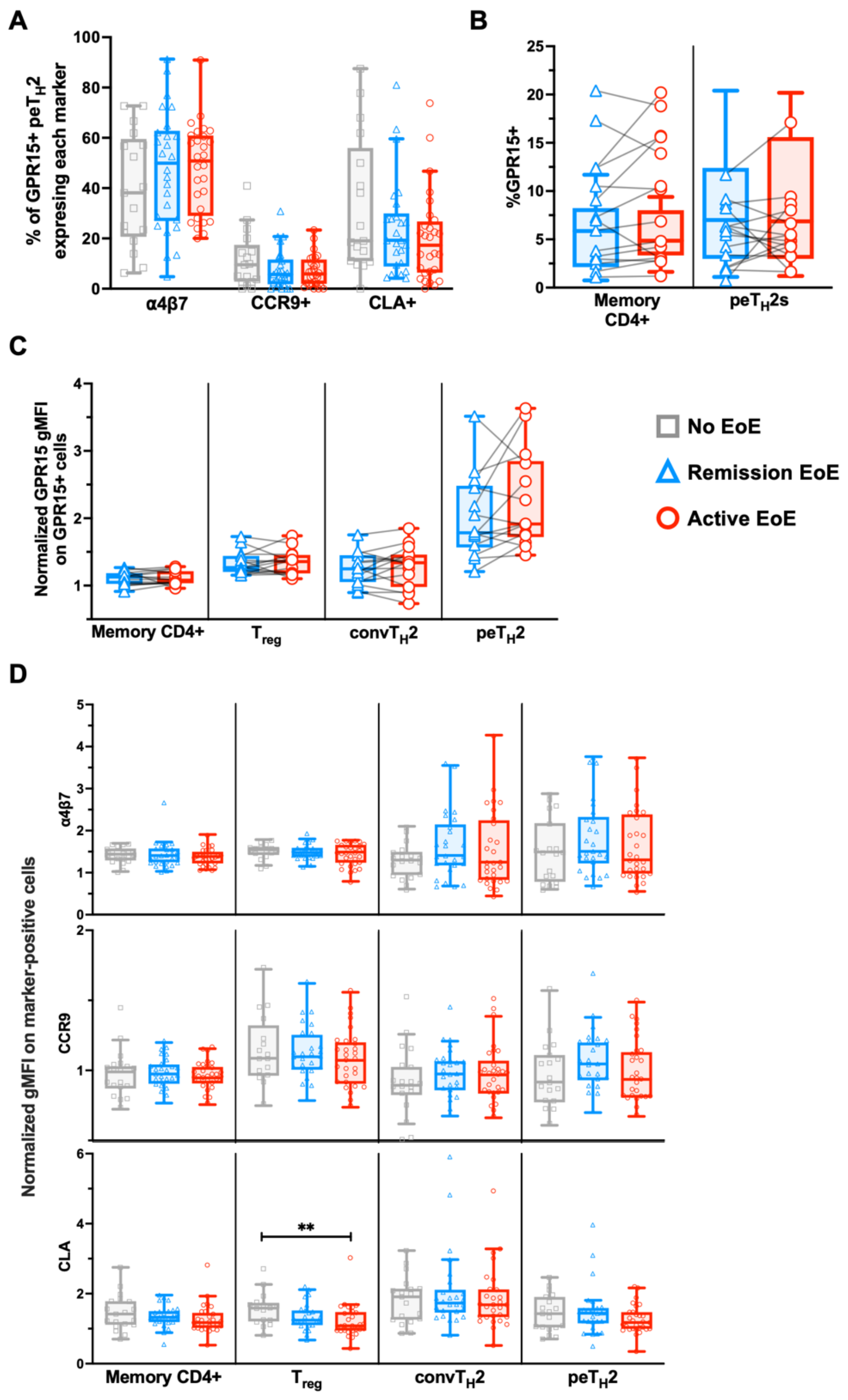


**Figure S1. GPR15 expression in paired samples and MFI of other markers. (A)** Percentage of GPR15+ peTh2 cells expressing other homing markers. **(B)** Percentage of peripheral blood memory CD4+ T cells and peTh2 cells expression GPR15 in the subset of paired samples. **(C)** Normalized gMFI of GPR15 on GPR15+ CD4+ T cell subsets in the subset of paired samples. MFI normalized to CD3+CD4-CD45RA-GPR15+ cells. **(D)** Normalized gMFI of other homing markers on marker-positive CD4+ T cell subsets. MFI normalized to CD3+CD4-CD45RA-marker+ cells. Paired samples were evaluated with Wilcoxon signed-rank tests. Differences between disease states were evaluated using Kruskal-Wallis tests followed by Wilcoxon rank-sum tests. **p<0.01 *peTh2, pathogenic effector Th2. EoE, eosinophilic esophagitis. convTh2, conventional Th2. gMFI, geometric mean fluorescence intensity. Treg, T regulatory.*

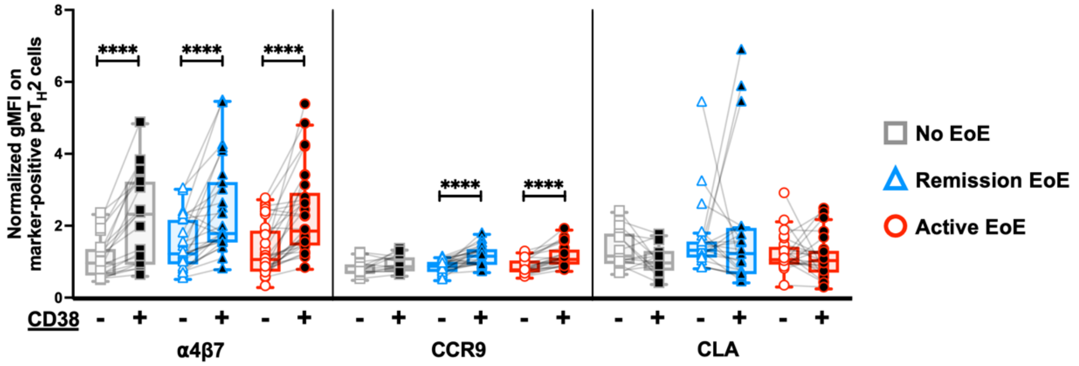


**Figure S2. CD38 expression by peripheral blood peTh2 cells and MFI of non-GPR15 migration markers.** Normalized gMFI of α4β7, CCR9, and CLA on marker-positive peT$_H$2s that were CD38+ or CD38-. MFI is normalized to CD3+CD4-CD45RA-marker+ cells. Lines connect cell populations in the same samples. Paired samples were evaluated with Wilcoxon signed-rank tests. ****$p<0.0001$. *peT$_H$2, pathogenic effector T$_H$2. EoE, eosinophilic esophagitis. gMFI, geometric mean fluorescence intensity.*

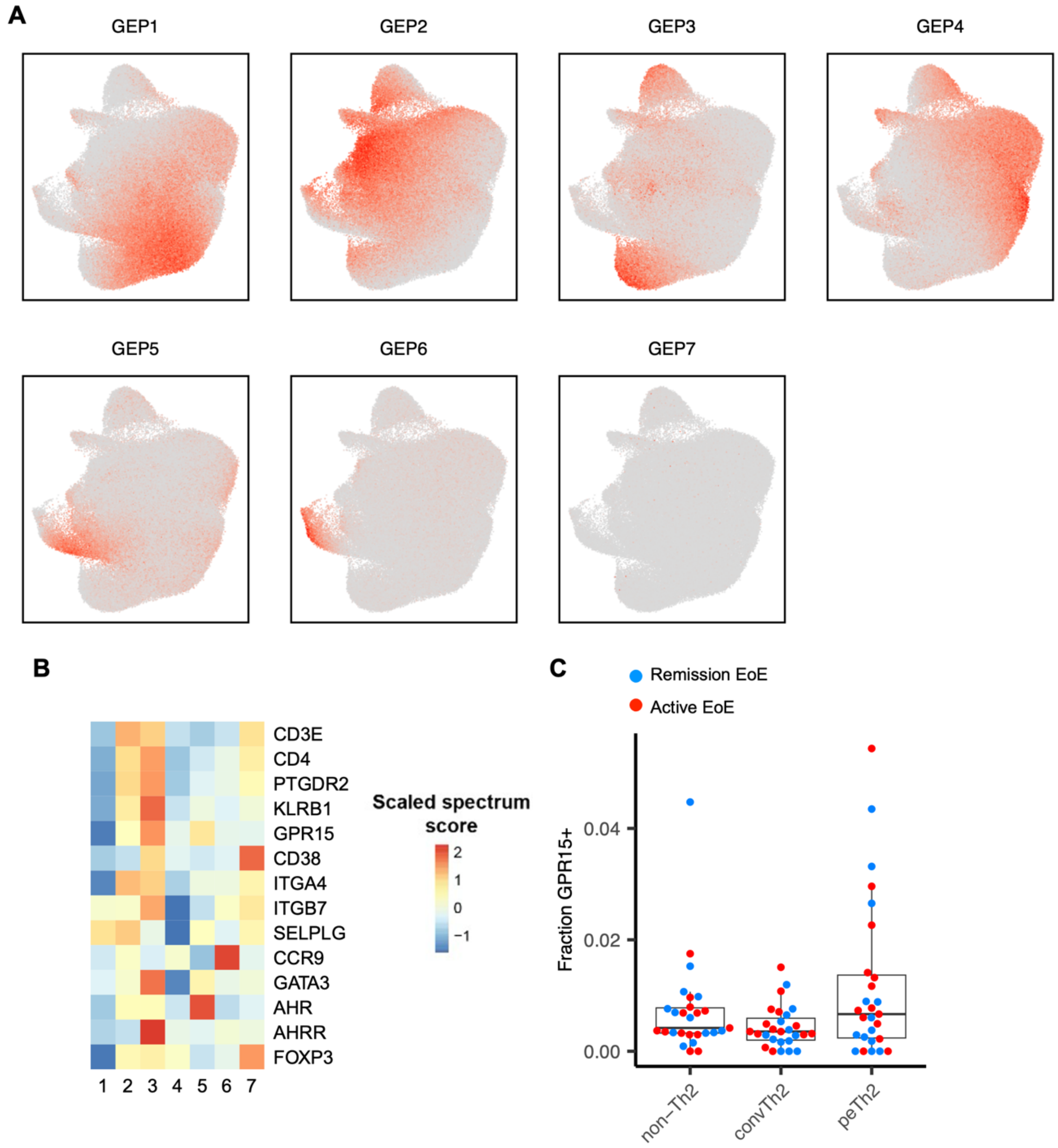


**Figure S3. Single cell supplemental data. (A)** Usage of each GEP for each cell overlaid on the UMAP visualization. **(B)** Scaled gene spectrum scores for genes that encode markers in flow panels in each gene expression program inferred using cNMF. **(C)** Fraction of non-Th2, convTh2, and peTh2 cells with GPR15 transcript detected in each patient. *GEP, gene expression program. cNMF, consensus negative matrix factorization. EoE, eosinophilic esophagitis. convTh2, conventional Th2. peTh2, pathogenic effector Th2.*

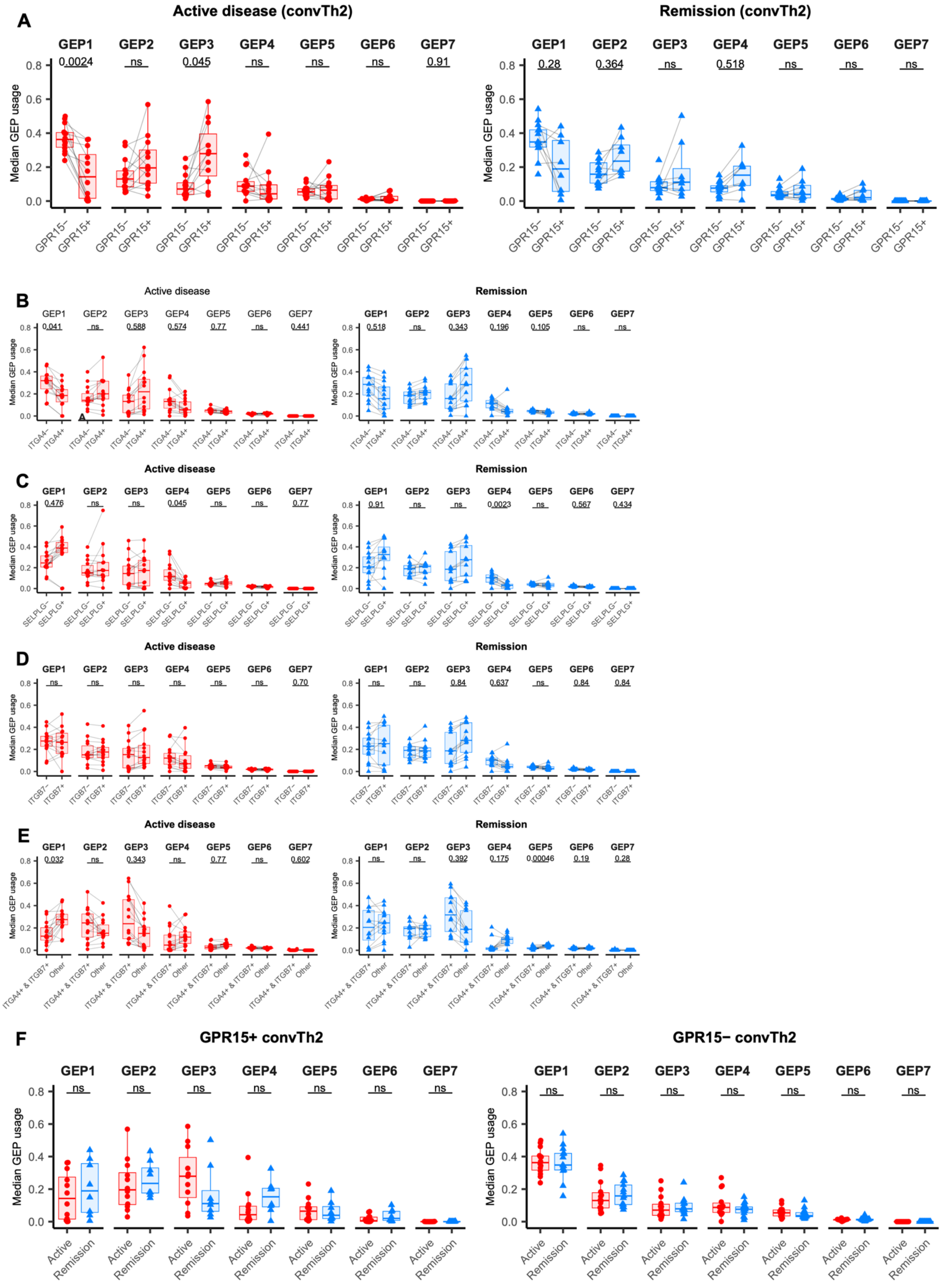


**Figure S4. GPR15+ convTh2 cells and peTh2 cells expressing α4β7 and CLA do not display the same gene expression patterns seen in GPR15+ peTh2 cells in EoE. (A)** Median normalized usage of each GEP among GPR15+ and GPR15- convTh2 cells from samples with active disease and remission. All samples shown have greater than one GPR15+ convTh2 cell. P-values are calculated with a two-sided t-test and are adjusted with Bonferroni correction. Median normalized usage of each GEP among peTh2 cells stratified by expression of: *ITGA4* **(B)**, *SELPLG* **(C)**, *ITGB7* **(D)**, or *ITGA4* and *ITGB7* co-expression **(E)** from samples active disease and remission. All samples shown have greater than one cell. P-values are calculated with a two-sided t-test and are adjusted with Bonferroni correction. **(F)** Median normalized usage of each GEP among GPR15+ and GPR15- convTh2 cells from samples with active disease and remission. All samples shown have greater than one GPR15+ convTh2 cell. P-values are calculated with a two-sided t-test and are adjusted with Bonferroni correction. *convTh2, conventional Th2. peTh2, pathogenic effector Th2. GEP, gene expression program. EoE, eosinophilic esophagitis.*

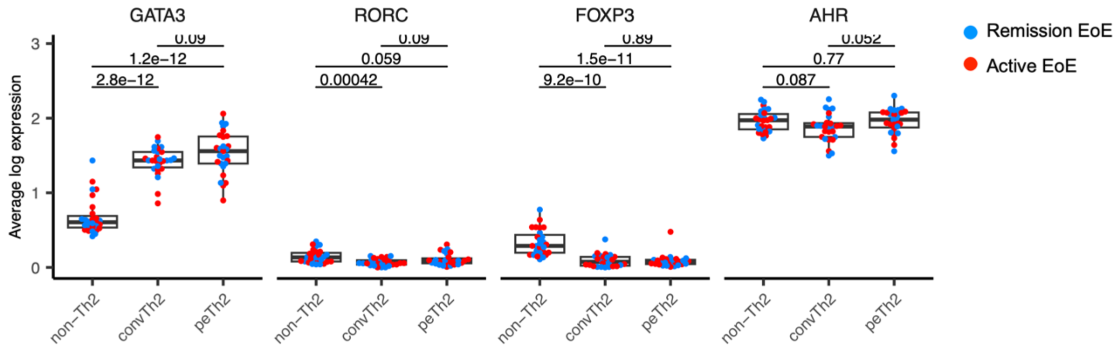


**Figure S5. Expression of key transcription factors in peripheral blood subsets in EoE.** Average pseudobulk expression of *GATA3, RORC, FOXP3,* and *AHR* among non-Th2, convTh2, and peTh2 cells from samples with active disease and remission. P-values are calculated with a two-sided t-test. *convTh2, conventional Th2. peTh2, pathogenic effector Th2.*

---

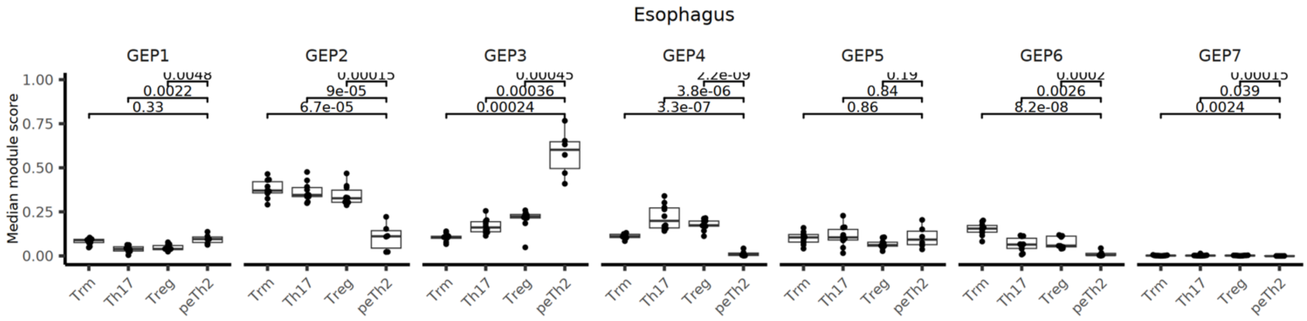


**Figure S6. GEP usage by esophageal T cells in EoE.** Median normalized usage of each GEP among esophageal CD8+ Trm, Th17, Treg, and peTh2 cells in EoE. P-values are calculated with a two-sided t-test and are adjusted with Bonferroni correction. *EoE, eosinophilic esophagitis. Trm, tissue resident memory. Treg, T regulatory. peTh2, pathogenic effector Th2.*

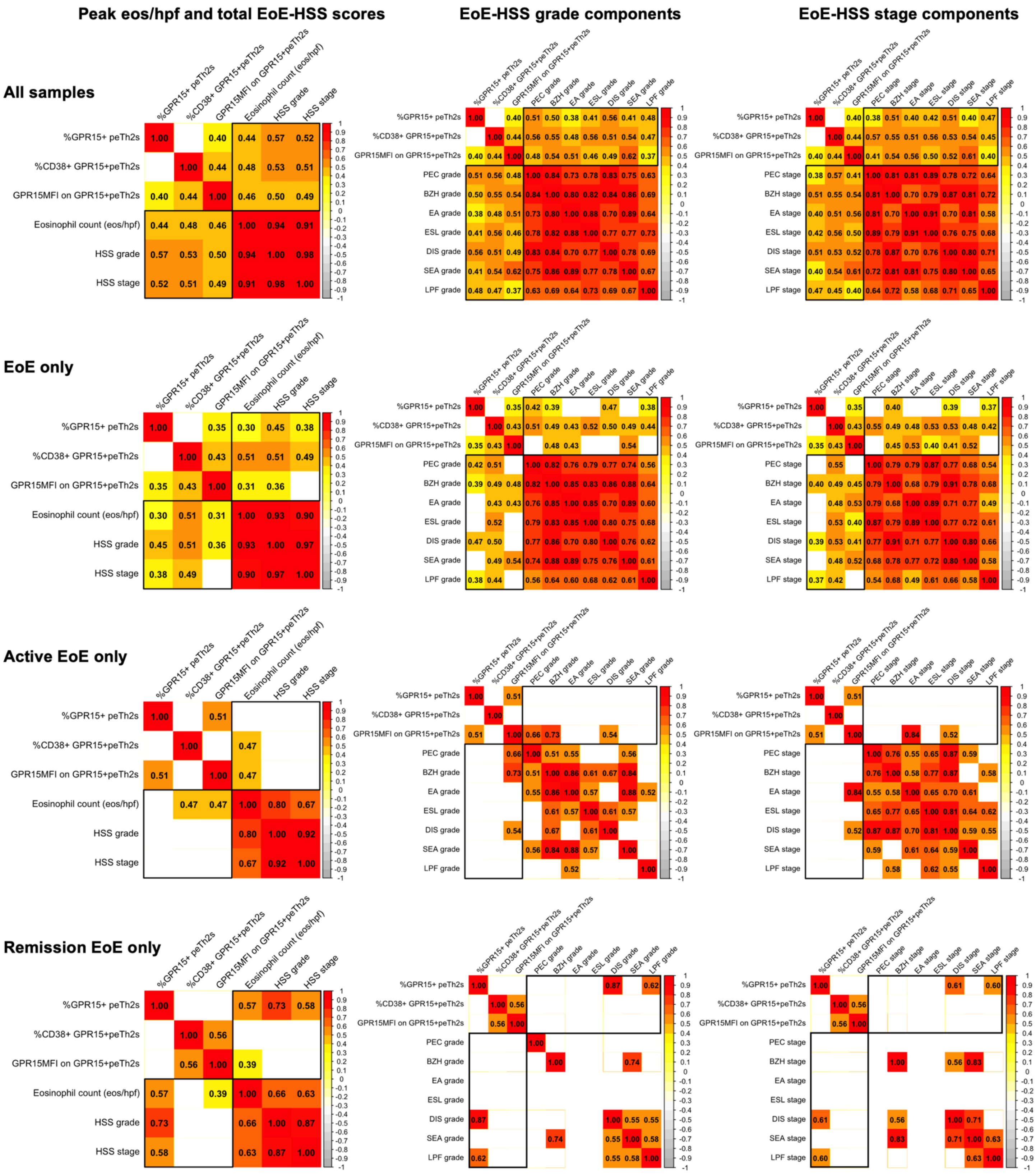


**Figure S7. Characteristics of peripheral blood peTh2 cells correlate with peak esophageal eosinophil count and EoE-HSS scores.** Correlation heatmaps for samples with paired peripheral blood flow cytometry data and clinical esophageal eosinophil count and/or research EoE-HSS data. Each row represents a subset of samples. Black outlines highlight correlations between peripheral blood and esophageal data. Correlation coefficients for Spearman correlations with P <0.05 shown. DEC scores are not shown; very few samples had non-zero scores. *peTh2, pathogenic effector Th2. EoE, eosinophilic esophagitis. EoE-HSS, EoE histologic*

*scoring system. Eos/hpf, eosinophils per high-power field. PEC, peak eosinophil count. BZH, basal cell hyperplasia. EA, eosinophilic abscesses. ESL, eosinophil surface layering. SEA, surface epithelial alteration. LPF, lamina propria fibrosis.*

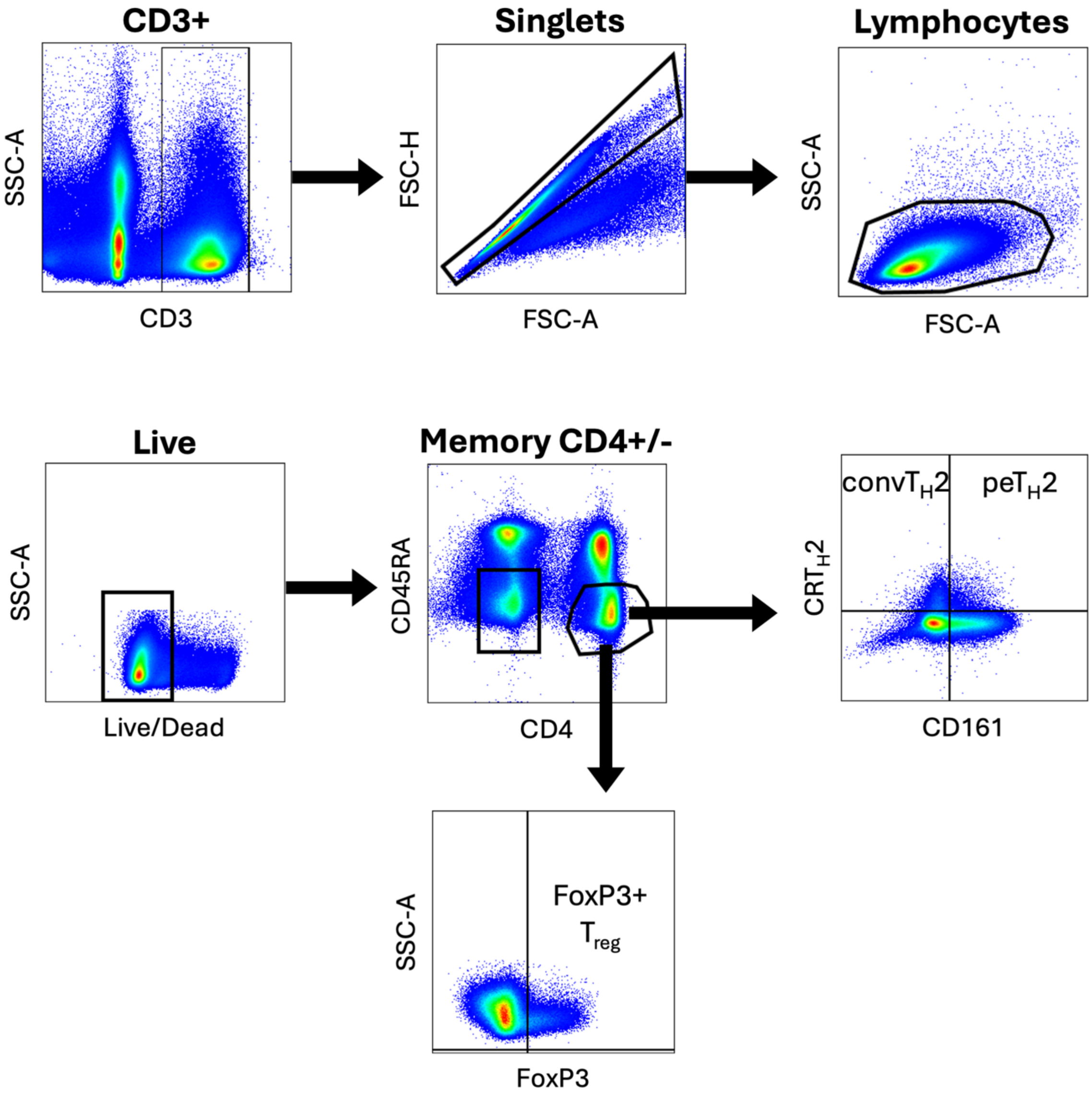


**Figure S8. Gating strategy for peripheral blood CD4+ populations examined in this report.** Memory CD4+ cells were defined as Live, singlet, lymphocytes that were CD3+CD4+CD45RA-. convTh2 cells were defined as CRTH2+CD161-. peTh2 cells were defined as CRTH2+CD161-. Tregs were defined as FoxP3+. *convTh2, conventional Th2. peTh2, pathogenic effector Th2. Treg, T regulatory.*

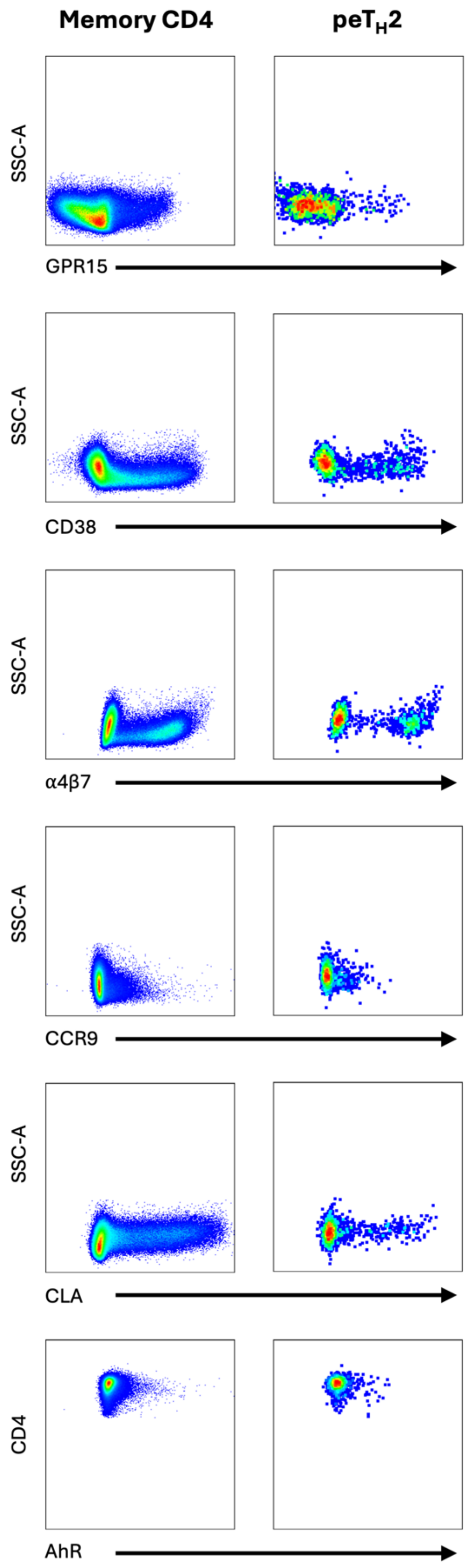


**Figure S9. Representative staining of key markers.** Flow cytometric plots of key markers on peripheral blood memory CD4+ cells (Live, CD3+CD4+CD45RA-) and peTh2 cells (Live, CD3+CD4+CD45RA-CRTh2+CD161+) from the same sample. *peTh2, pathogenic effector Th2.*
