## Supplemental tables for "GPR15 and CD38 define a subset of peripheral blood pathogenic effector Th2 cells associated with active eosinophilic esophagitis"

**Table S1. Summary demographic characteristics of subjects included in peripheral blood flow cytometry data.**

| All samples | EoE, active ($n$* = 29) | EoE, remission ($n$ = 26) | Not EoE ($n$ = 19) | $p^{\dagger}$ |
|---|---|---|---|---|
| Age (years), median (IQR) | 11 (10) | 16 (8) | 6 (10) | 0.14 |
| Male, *n* (%) | 22 (76) | 22 (85) | 10 (52) | 0.062 |
| Race, *n* (%) | | | | |
| White | 28 (97) | 25 (96) | 18 (95) | 1 |
| Black/African American | 0 (0) | 0 (0) | 1 (5) | 1 |
| Asian | 0 (0) | 0 (0) | 1 (5) | 1 |
| Ethnicity, *n* (%) | | | | |
| Hispanic/Latino | 1 (3) | 0 (0) | 2 (11) | 0.26 |
| Allergic comorbidities, *n* (%) | | | | |
| Asthma | 12 (41) | 13 (50) | 6 (32) | 0.46 |
| Allergic rhinitis | 8 (28) | 6 (23) | 11 (58) | **0.047** |
| Eczema | 10 (29) | 8 (31) | 7 (37) | 0.95 |
| IgE-mediated food allergy | 10 (34) | 8 (30) | 12 (63) | 0.072 |
| FPIES | 0 (0) | 0 (0) | 1 (5) | 0.26 |
| Drug allergy | 5 (17) | 4 (15) | 2 (11) | 0.92 |
| Family history, *n* (%) | | | | |
| Of atopy‡ | 19 (66) | 15 (58) | 14 (74) | 0.58 |
| Of EoE | 6 (21) | 1 (4) | 1 (5) | 0.10 |

*EoE, eosinophilic esophagitis. IQR, interquartile range. IgE, immunoglobulin E. FPIES, food-protein-induced enterocolitis syndrome.*

*Longitudinal samples were considered independent. Individual characteristics are included in **Tables S3-4**.

†Fisher's exact test for proportions, Kruskal-Wallis test for continuous variables.

‡Atopy was defined as asthma, eczema, IgE-mediated food allergy, allergic rhinitis.

**Table S2. Summary endoscopic and histological characteristics of subjects included in peripheral blood flow cytometry data who underwent endoscopy.**

| Samples with associated endoscopy | | EoE, active ($n$* = 29) | EoE, remission ($n$ = 26) | Not EoE ($n$ = 11) | $p^{\dagger}$ |
|---|---|---|---|---|---|
| Eosinophil count (eos/hpf)‡, median (IQR, maximum) | | | | | |
| Peak | | 46 (30, 100) | 1 (5, 10) | 0 (0, 1) | **<0.0001** |
| Distal esophagus | | 35 (40, 90) | 0.25 (3.8, 10) | 0 (0, 1) | **<0.0001** |
| Middle esophagus | | 20 (45, 100) | 0 (2.5, 9) | 0 (0, 1) | **<0.0001** |
| Proximal esophagus | | 25 (40, 75) | 0 (0.38, 3) | 0 (0, 0) | **<0.0001** |
| EREFS | | | | | |
| Scored, *n* (%) | | 25 (86) | 22 (85) | 11 (100) | |
| If scored, median (IQR, maximum) | | | | | |
| Total | | 3 (2, 7) | 0.5 (2, 4) | 0, (0, 4) | **<0.0001** |
| Edema | | 1 (1, 1) | 0 (0, 1) | 0 (0, 0) | **0.0002** |
| Rings | | 0 (1, 2) | 0 (1, 2) | 0 (0, 1) | 0.37 |
| Exudate | | 1 (1, 2) | 0 (1, 1) | 0 (0, 1) | **<0.0001** |
| Furrows | | 1 (0, 1) | 0 (1,1) | 0 (0,1) | **0.0001** |
| Stricture | | 0 (0,1) | 0 (0, 0) | 0 (0, 0) | 0.26 |
| Peak EoE-HSS | | | | | |
| Scored, *n* (%) | | 16 (55) | 15 (58) | 11 (100) | |
| If scored, median (IQR, maximum) | | | | | |
| Overall | G | 0.56 (0.23, 0.88) | 0.042 (0.085, 0.33) | 0 (0.021, 0.083) | **<0.0001** |
| | S | 0.58 (0.26, 0.67) | 0.04 (0.06, 0.42) | 0 (0.02, 0.08) | **<0.0001** |
| Eosinophilic inflammation (EI) | G | 2 (1 , 3) | 1 (1, 1) | 0 (0 ,1) | **<0.0001** |
| | S | 3 (0.25, 3) | 0 (0, 0) | 0 (0, 0) | **<0.0001** |
| Basal zone hyperplasia (BZH) | G | 3 (1, 3) | 0 (1, 2) | 0 (0, 1) | **<0.0001** |
| | S | 3 (0.25, 3) | 0 (1, 3) | 0 (0 ,1) | **<0.0001** |
| Eosinophil abscess (EA) | G | 1 (1, 3) | 0 (0, 0) | 0 (0, 0) | **<0.0001** |
| | S | 1 (2, 2) | 0 (0, 0) | 0 (0, 0) | **<0.0001** |
| Eosinophil surface layering (SL) | G | 1.5 (1, 3) | 0 (0, 0) | 0 (0, 0) | **<0.0001** |
| | S | 2 (1, 3) | 0 (0, 0) | 0 (0, 0) | **<0.0001** |
| Dilated intracellular spaces (DIS) | G | 3 (0.25, 3) | 1 (2, 2) | 0 (0, 1) | **<0.0001** |
| | S | 3 (1, 3) | 1 (1.5, 3) | 0 (0, 1) | **<0.0001** |
| Surface epithelial alteration (SEA) | G | 2 (1.3, 3) | 0 (0, 1) | 0 (0, 0) | **<0.0001** |
| | S | 1 (0.5, 3) | 0 (0 ,1) | 0 (0, 0) | **<0.0001** |
| Dyskeratotic epithelial cells (DEC) | G | | | | 0.44 |
| | S | 0 (0, 0) | 0 (0,0) | 0 (0, 0) | 1 |
| Lamina propria fibrosis (LPF) | G | 2 (2.3, 3) | 0 (0, 2) | 0 (0, 0) | **<0.0001** |
| | S | 1 (1.3, 3) | 0 (0, 3) | 0 (0, 0) | **0.00032** |

*EoE, eosinophilic esophagitis. Eos/hpf, eosinophils per high-power field. EREFS, endoscopic reference score.*

*Longitudinal samples were considered independent. Individual characteristics are included in **Tables S3-4**.

†Kruskal-Wallis test.

‡samples reported clinically as “>40” and were unable to be re-evaluated are imputed as 60 eos/hpf. samples reported as “rare” are imputed as 0.5 eos/hpf.

**Table S3. Individual characteristics of subjects undergoing endoscopy included in peripheral blood flow cytometry data.**

| Subject ID | Age (yrs) | Sex | Race | Ethnicity | Allergic comorbidities | Family History | | EoE trigger(s) | Eosinophil count (eos/hpf) | | | EREFS | EoE-HSS (peak total) | | Group |
|---|---|---|---|---|---|---|---|---|---|---|---|---|---|---|---|
| | | | | | | Atopy* | EoE | | D | M | P | | Grade | Stage | |
| 1 | 16 | M | W | N | A, IgE-FA | N | N | tree nut, potato, turkey, corn | 1 | 0 | 0 | 0/0/0/0/0 = 0 | 0.04 | 0.04 | R |
| | | | | | | | | | 18 | 1 | 1 | 0/0/0/0/0 = 0 | 0.17 | 0.17 | A |
| | 17 | | | | | | | | rare | 0 | rare | 0/0/0/0/0 = 0 | NS | NS | R |
| 2 | 16 | M | W | N | AR, AD, DA | Y | N | milk, soy, egg | 0 | 0 | 0 | NS | 0.08 | 0.04 | R |
| | 17 | | | | | | | | 24 | 20 | 12 | NS | 0.29 | 0.38 | A |
| 14 | 20 | M | W | N | AR | N | N | milk, beef | 75 | 100 | 50 | NS | 0.88 | 0.67 | A |
| | 21 | | | | | | | | 5 | 0 | 0 | 0/0/0/0/0 = 0 | 0.08 | 0.04 | R |
| 72 | 15 | M | W | N | A | Y | N | egg | 5 | 12 | 25 | 1/2/1/1/0 = 5 | NS | NS | A |
| | 16 | | | | | | | | 10 | 3 | 0 | 0/0/1/1/0 = 2 | NS | NS | R |
| | 18 | | | | | | | | 15 | 40 | 30 | 1/2/2/1/1 = 7 | NS | NS | A |
| | 22 | | | | | | | | 7 | 0 | 0 | 0,2,0,0,0 = 2 | NS | NS | R |
| 131 | 27 | F | W | N | AD | Y | N | milk | 0 | 8 | 0 | NS | 0.29 | 0.29 | R |
| | 30 | | | | | | | | 30 | 5 | 45 | 0/0/2/1/0 = 3 | 0.42 | 0.29 | A |
| 175 | 6 | M | W | N | A, AR AD, IgE-FA | Y | N | meat | >40 | >40 | 35 | NS | NS | NS | A |
| | 7 | | | | | | | | 0 | 5 | rare | 0/0/1/1/0 = 2 | NS | NS | R |
| 288 | 3 | M | W, A | N | A, AD, DA, IgE-FA | Y | N | soy | 9 | 18 | 9 | NS | NS | NS | A |
| | 4 | | | | | | | | 0 | 4 | 0 | 0/0/1/1/0 = 2 | NS | NS | R |
| 293 | 3 | M | A | N | DA | N | N | milk | 0 | 3 | VR | NS | 0 | 3 | R |
| | | | | | | | | | >40 | >40 | >40 | NS | >40 | >40 | A |
| 318 | 22 | M | W | N | A, AR | N | N | milk | 7 | 1 | 0 | 0/1/1/0/0 = 2 | NS | NS | R |
| | 23 | | | | | | | | 20 | 35 | 5 | 0/1/1/1/0 = 3 | NS | NS | A |
| 348 | 12 | F | W | N | *none* | N | N | milk, wheat | 46 | 3 | 45 | 0/0/1/1/0 = 2 | 0.46 | 0.42 | A |
| | 18 | | | | | | | | 0 | 0 | 0 | 0/0/0/0/0 = 0 | 0.00 | 0.00 | R |
| 362 | 5 | M | W | N | *none* | N | Y | milk | 10 | 6 | 3 | NS | NS | NS | R |
| | 6 | | | | | | | | 28 | rare | 3 | NS | NS | NS | A |
| 448 | 3 | M | W | N | *none* | Y | N | milk, egg, or wheat | 4 | 9 | 3 | 0/0/0/0/0 = 0 | 0.33 | 0.42 | R |
| | 8 | | | | IgE-FA | | | | 40 | 30 | 25 | 1/0/1/1/0 = 3 | 0.58 | 0.63 | A |
| 6425 | 17 | F | W | N | A, IgE-FA | Y | N | milk | 35 | 8 | 3 | 0/2/2/1/0 = 5 | 0.33 | 0.33 | A |
| | | | | | | | | | 0 | 0 | 0 | 0/1/1/1/0 = 3 | 0.00 | 0.00 | R |
| 6577 | 4 | F | W | N | A, IgE-FA | Y | N | milk | 10 | 50 | 45 | 1/0/1/1/0 = 3 | 0.67 | 0.67 | A |
| | | | | | | | | | rare | rare | 0 | 0/0/0/0/0 = 0 | NS | NS | R |

| | | | | | | | | | | | | | | | |
|---|---|---|---|---|---|---|---|---|---|---|---|---|---|---|---|
| 8010 | 6 | M | W | N | *none* | N | Y | milk | 20 | 15 | 25 | 1/0/2/1/0 = 4 | NS | NS | A |
| | 7 | | | | | | | | >40 | 3 | 0 | 1/0/1/1/0 = 3 | NS | NS | A |
| 291 | 7 | M | W | OSHL | A, AR | Y | N | *unknown* | >40 | >40 | 35 | 0/0/1/1/0 = 2 | NS | NS | A |
| 0587 | 4 | F | W | N | A, AD, IgE-FA, DA | N | Y | milk | 55 | 17 | 75 | 1/0/1/1/0 = 3 | 0.63 | 0.58 | A |
| 1582 | 16 | M | W | N | AD | Y | Y | *unknown* | 90 | 100 | 45 | 1/2/1/1/0 = 5 | 0.71 | 0.67 | A |
| 2137 | 16 | M | W | N | *none* | Y | N | *unknown* | 26 | 35 | 25 | 1/0/2/1/0 = 4 | 0.58 | 0.63 | A |
| 2560 | 18 | F | W | N | *none* | Y | N | *unknown* | 20 | 65 | 45 | 0/2/2/1/0 = 5 | 0.54 | 0.63 | A |
| 2737 | 6 | F | W | N | A, IgE-FA, DA | Y | N | *unknown* | 65 | 42 | 58 | 1/1/1/1/0 = 4 | 0.54 | 0.58 | A |
| 3621 | 14 | M | W | N | AR, E, IgE-FA | Y | N | milk | 25 | 2 | 1 | 0/1/0/1/0 = 2 | 0.25 | 0.17 | A |
| 4709 | 9 | M | W | N | AD, AR | Y | Y | Salmon, nuts/seeds | >40 | 5 | 8 | 0,1,1,1,0 = 3 | NS | NS | A |
| 5185 | 10 | M | W | N | *none* | Y | N | milk | 30 | 0 | 0 | 0/0/0/0/0 = 0 | NS | NS | A |
| 7960 | 14 | M | W | N | AR | Y | N | *unknown* | 55 | 15 | 21 | 0/1/2/1/0 = 4 | 0.63 | 0.63 | A |
| 8173 | 10 | M | W | N | A, AD, IgE-FA | Y | N | *unknown* | 90 | 30 | 3 | 0/1/2/1/0 = 4 | 0.58 | 0.58 | A |
| 9809 | 11 | M | W | N | AD | N | N | *unknown* | >40 | >40 | >40 | 1/0/2/1/1 = 5 | NS | NS | A |
| 424 | 18 | M | W | N | A, AD, IgE-FA | Y | N | milk | 0 | 0 | 0 | NS | 0.13 | 0.08 | R |
| 2542 | 11 | M | W | N | A, AR, AD | Y | N | milk | 0 | 1 | 0 | 0/0/0/0/0 = 0 | 0.04 | 0.00 | R |
| 3295 | 16 | M | W | N | *none* | N | N | milk | 0 | 0 | 0 | 0/0/0/0/0 = 0 | 0.04 | 0.04 | R |
| 3594 | 10 | M | W | N | DA | Y | N | milk | rare | 0 | 0 | 0/0/1/0/0 = 0 | NS | NS | R |
| 4772 | 17 | M | W | N | AR, AD | Y | N | milk | 0 | 0 | 0 | 0/1/0/1/0 = 2 | 0.00 | 0.00 | R |
| 5614 | 16 | M | W | N | A, AD, IgE-FA | Y | N | milk | 3 | 0 | rare | 0/0/0/0/0 = 0 | NS | NS | R |
| 6300 | 12 | M | W | N | A | N | N | milk | 1 | 0 | 0 | 1/1/1/1/0 = 4 | 0.08 | 0.04 | R |
| 7064 | 18 | M | W | N | *none* | N | N | milk or chicken | 0 | 0 | 0 | 0/0/0/0/0 = 0 | 0.00 | 0.00 | R |
| 7830 | 24 | M | W | N | *none* | N | N | milk | 4 | 0 | 0 | 0/1/0/0/0 = 1 | 0.21 | 0.29 | R |
| 9056 | 11 | M | W | N | *none* | Y | N | milk | 0 | 0 | 1 | 0/1/0/0/0 = 1 | 0.04 | 0.08 | R |
| 0881 | 17 | M | W | N | AR, AD, IgE-FA,DA | Y | N | -- | 0 | 0 | 0 | 0/0/0/0/0 = 0 | 0.00 | 0.00 | N |
| 2646 | 12 | M | W | PR | *none* | Y | N | -- | 0 | 0 | 0 | 0/0/0/0/0 = 0 | 0.00 | 0.00 | N |
| 2855 | 20 | F | W | N | IgE-FA | N | Y | -- | 0 | 0 | 0 | 0/0/0/0/0 = 0 | 0.00 | 0.00 | N |
| 2905 | 4 | M | W | OHSL | AR | Y | N | -- | 0 | 1 | 0 | 0/0/0/0/0 = 0 | 0.08 | 0.08 | N |
| 3141 | 19 | F | W | N | *none* | Y | N | -- | 0 | 0 | 0 | 0/1/1/1/0 = 3 | 0.04 | 0.04 | N |
| 5137 | 5 | M | W | N | *none* | Y | N | -- | 0 | 0 | 0 | 0/0/0/0/0 = 0 | 0.00 | 0.00 | N |
| 5667 | 6 | M | W | N | A, AR, FPIES | Y | N | -- | 0 | 0 | 0 | NS | 0.00 | 0.00 | N |
| 6264 | 11 | F | W | N | *none* | Y | N | -- | 1 | 0 | 0 | 0/0/0/0/0 = 0 | 0.08 | 0.04 | N |
| 7265 | 6 | F | W | N | *none* | Y | N | -- | 0 | 0 | 0 | NS | 0.00 | 0.00 | N |

| | | | | | | | | | | | | | | | |
|---|---|---|---|---|---|---|---|---|---|---|---|---|---|---|---|
| 7423 | 3 | F | W | N | AD, IgE-FA, DA | Y | N | -- | 0 | 0 | 0 | 0/0/0/0/0 = 0 | 0.00 | 0.00 | N |
| 7565 | 18 | F | W | N | IgE-FA | Y | N | -- | 0 | 0 | 0 | 0/1/1/1/0 = 3 | 0.00 | 0.00 | N |

*eos/hpf, eosinophils per high-power field. D, distal. M, mid. P, proximal. EREFS, endoscopic reference score. EoE-HSS, eosinophilic esophagitis histologic scoring system (reported where available).*

*Atopy in family history defined as asthma, eczema, allergic rhinitis

Key:

- Sex: M = male, F = female
- Race: W = white, A = asian
- Ethnicity: N = not Hispanic/Latino, OSHL = other Spanish/Hispanic/Latino, PR = Puerto Rican
- Allergic comorbidities: A = asthma, IgE-FA = IgE-mediated food allergy, AR = allergic rhinitis, AD = atopic dermatitis/eczema, DA = drug allergy, FPIES = food-protein-induced enterocolitis syndrome
- Eosinophil count: VR = “very rare”
- EREFS: scores reported as edema/rings/exudates/furrows/strictures = total
- EREFS or HSS: NS = not scored
- Group: A = active EoE, R = EoE in remission after diet therapy, N = not EoE

**Table S4. Individual characteristics of subjects enrolled in IgE-mediated food allergy studies included in peripheral blood flow cytometry data.**

| Subject ID | Selected from | Age (yrs) | Sex | Race | Ethnicity | Allergic comorbidities | Family History | | Food allergies | Previous reaction(s) | Food allergy testing† or positive challenge during clinical trial |
|---|---|---|---|---|---|---|---|---|---|---|---|
| | | | | | | | Atopy* | EoE | | | |
| 33 | Biorepository | 5 | M | W | N | A, AR | N | N | Peanut | C, T | IgE ($kU_A$/L): peanut 16.7, Ara h 2 11.8<br>SPT (mm) 6 months after sample: 9/25 |
| 47 | Biorepository | 7 | M | W | N | A, AR | N | N | Peanut | C, GI | IgE ($kU_A$/L): peanut 97.8, Ara h 2 >100<br>SPT (mm): 16/32 |
| 89 | Biorepository | 6 | M | W | N | A, AR, AD | Y | N | Egg | C, T, GI | IgE ($kU_A$/L): egg white 1.14, ovomucoid 0.89<br>SPT (mm): egg white 10/27, egg yolk 6/20 |
| | | | | | | | | | Milk | C, R | IgE ($kU_A$/L): milk 4.39, casein 1.79, whey 5.30<br>SPT (mm): milk 20/25, casein 6/9 |
| | | | | | | | | | Peanut | C, R | IgE ($kU_A$/L): peanut 87.7, Ara h 2 78.4<br>SPT (mm): 18/28 |
| | | | | | | | | | Sunflower seed | C, M | IgE ($kU_A$/L): 5.84<br>SPT (mm): 10/24 |
| 102 | Biorepository | 3 | M | W | N | A, AR, AD | Y | N | Egg | C, R | IgE ($kU_A$/L): egg white 0.44, ovomucoid 0.14<br>SPT (mm) 2 years before sample: egg white 3/11, egg yolk 0/7 |
| | | | | | | | | | Peanut | C | IgE ($kU_A$/L): peanut 31.2, Ara h 2 23.7<br>SPT (mm) 10 months after sample: 43/50 |
| | | | | | | | | | Shellfish | C | IgE ($kU_A$/L): clam 6.26, oyster 0.82, scallop 4.03, crab 20.3, lobster 19.3, shrimp 20.1, squid 0.84.<br>SPT not available |
| | | | | | | | | | Beef | C, M | IgE ($kU_A$/L): 1.11<br>SPT (mm) 10 months after sample: 35/50 |
| | | | | | | | | | Lamb | C | IgE ($kU_A$/L): 1.04<br>SPT (mm) 10 months after sample: 25/50 |
| 105 | Biorepository | 12 | F | W | N | AR, AD | N | N | Egg | C, GI, M | IgE ($kU_A$/L): egg white 0.39<br>SPT (mm): egg white 0/0 |
| | | | | | | | | | Peanut | U | IgE ($kU_A$/L): peanut >100, Ara h 2 67.3.<br>SPT (mm) 2 years before sample: 25/29 |
| 6 | Clinical trial | 27 | F | A, BAA | N | AR, AD | N | N | Tree nut | E, M, T | Positive DBPFC to hazelnut |
| 13 | Clinical trial | 6 | M | W | N | A, AR, AD | Y | N | Peanut | C, GI, M, R, T | Positive DBPFC to peanut |
| | | | | | | | | | Tree nut | C | |
| | | | | | | | | | Shellfish | U | |
| 15 | Clinical trial | 5 | F | W | N | AR | Y | N | Peanut | GI, T | Positive DBPFC to peanut |
| | | | | | | | | | Tree nut | C, GI | |

*SPT, skin prick test. mm, millimeters. DBPFC, double-blind, placebo-controlled food challenge.*

*Atopy in family history defined as asthma, eczema, allergic rhinitis. IgE, immunoglobulin E. †testing within 2 weeks of sample collection unless otherwise noted.

Key:

- Sex: M = male, F = female
- Race: W = white, A = Asian, BAA = Black/African American
- Ethnicity: N = not Hispanic/Latino
- Allergic comorbidities: A = asthma, AR = allergic rhinitis, AD = atopic dermatitis/eczema
- Previous reactions: C = cutaneous; T = throat clearing, throat hoarseness, or cough; R = respiratory; GI = gastrointestinal; M = mouth or throat itching; E = eye itching, eye redness, eye discharge; perioral hives; U = unknown

**Table S5. Fluorescently-labeled antibody panel used in peripheral blood flow cytometry data.**

| Marker | Fluorochrome | Clone | Manufacturer |
|---|---|---|---|
| CD3 | BUV395 | UCHT1 | BD Biosciences |
| Live/Dead Fixable Blue | | - | Invitrogen |
| CD4 | BUV496 | RPA-T4 | BD Biosciences |
| *CD49d** | *BUV661* | *9F10* | *BD Biosciences* |
| *CCR7* | *BUV737* | *2-L1-4* | *BD Biosciences* |
| CCR9 | BV421 | L053E8 | Biolegend |
| CD161 | BV480 | HP-3G10 | BD Biosciences |
| CD45RA | BV650 | HI100 | BD Biosciences |
| *OX40* | *BV711* | *Ber-ACT35* | *Biolegend* |
| *CD27* | *BV750* | *O323* | *Biolegend* |
| α4β7 | AlexaFluor 488 | Hu117 | R&D Systems |
| *CD8A* | *BB700* | *OKT8* | *BD Biosciences* |
| *CD127* | *PerCP-eFluor710* | *eBioRDR5* | *eBioscience* |
| *RORγ-t*† | *PE* | *AFKJS-9* | *eBioscience* |
| AhR | PE-CF594 | T49-550 | BD Biosciences |
| CRTh2 | PE-Cy5 | BM16 | eBioscience |
| *CD25* | *PE-Fire700* | *M-A251* | *Biolegend* |
| *Tbet* | *PE-Cy7* | *4B10* | *Biolegend* |
| GPR15 | APC | SA302A10 | Biolegend |
| FoxP3 | efluor 660 | 236A/E7 | eBioscience |
| CD38 | APC-R700 | HB7 | BD Biosciences |
| CLA | APC-Vio770 | HECA-452 | Miltenyi |

*Analysis of markers in *italics* is not included in this report.
†Three samples in the "not food allergy" group were not subjected to intracellular staining (RORγ-t, AhR Tbet, FoxP3).

**Table S6. Individual characteristics of previously unpublished subjects included in peripheral blood single-cell RNA sequencing data.**

| Subj ID | Age (yrs) | Sex | Race | Ethnicity | Allergic comorbidities | Family History | | EoE trigger(s) | Eosinophil count (eos/hpf) | | | EREFS | Group |
|---|---|---|---|---|---|---|---|---|---|---|---|---|---|
| | | | | | | Atopy* | EoE | | D | M | P | | |
| 215 | 10 | F | W | N | none | Y | N | milk | 3 | >40 | >40 | NS | A |
| | 11 | | | | | | | | 0 | 0 | 0 | NS | R |
| | 18 | | | | | | | | 0 | 0 | 0 | NS | R |
| 417 | 12 | M | W | N | AD, DA | Y | N | milk | VR | VR | 0 | 0,1,0,1,0 = 2 | R |
| | 13 | | | | | | | | 35 | 35 | 0 | 0,1,2,1,0 = 4 | A |
| | 17 | | | | | | | | >40 | 12 | 0 | 0,1,2,0,0 = 3 | A |
| 433 | 6 | M | W | N | DA | Y | N | milk | 5 | <3 | VR | 0,0,0,1,0 = 1 | R |
| | 11 | | | | | | | | VR | VR | 0 | NS | R |
| 343 | 8 | M | W | N | A, AD, IgE-FA | Y | N | milk | VR | 0 | NA | 0,0,1,1,0 = 2 | R |
| | 14 | | | | A, AD, AR, IgE-FA | | | | 34 | 0 | 0 | 0,1,1,1,0 =3 | A |
| 8901 | 16 | F | W | N | none | Y | N | milk | >40 | 3 | 10 | NS | A |
| | | | | | | | | | 0 | 4 | 0 | 0,0,1,1,0 = 2 | R |
| 1788 | 5 | M | W | N | *none* | Y | Y | milk | 0 | 0 | 0 | NS | R |
| 0411 | 10 | F | W | N | *none* | Y | Y | milk | 15 | 20 | 28 | 1,0,1,1,0 = 3 | A |
| 440 | 15 | M | W | N | A, AD, AR | Y | N | milk | 0 | VR | 0 | NS | R |
| 478 | 21 | M | W | N | AR | N | N | milk, egg, legume | >40 | 0 | 5 | 0,2,2,1,0 = 5 | A |
| 489 | 14 | M | W | N | AD, AR | N | N | milk | 38 | 37 | VR | 0,1,2,0,0 = 3 | A |
| 1052 | 12 | M | W, BAA | OSHL | A, IgE-FA | Y | N | *unknown* | 35 | >40 | >40 | 1,0,2,1,0 = 4 | A |
| 1112 | 16 | F | W | N | AR, DA | Y | Y | *unknown* | >40 | >40 | 30 | 1,1,1,0,0 = 3 | A |
| 6398 | 9 | F | W | N | AR | Y | N | milk | 4 | 5 | 3 | 0,0,0,1,0 = 1 | R |
| 7687 | 8 | M | W | N | A | Y | N | milk | 0 | 0 | 0 | NS | R |

*eos/hpf, eosinophils per high-power field. D, distal. M, mid. P, proximal. EREFS, endoscopic reference score.*
*Atopy in family history defined as asthma, eczema, allergic rhinitis.
Key:

- Sex: M = male, F = female
- Race: W = white, BAA = Black/African American
- Ethnicity: N = not Hispanic/Latino, OSHL = other Spanish/Hispanic/Latino, PR = Puerto Rican
- Allergic comorbidities: A = asthma, IgE-FA = IgE-mediated food allergy, AR = allergic rhinitis, AD = atopic dermatitis/eczema, DA = drug allergy, FPIES = food-protein-induced enterocolitis syndrome,
- Eosinophil count (eos/hpf): VR = “very rare”
- EREFS: scores reported as edema/rings/exudates/furrows/strictures = total, NS = not scored
- Group: A = active EoE, R = EoE in remission after diet therapy, N = not EoE

**Table S7. Individual characteristics of subjects included in esophagus flow cytometry data.**

| Subject ID | Age (yrs) | Sex | Race | Ethnicity | Allergic comorbidities | Family History | | EoE trigger(s) | Eosinophil count (eos/hpf) | | | EREFS | Group |
|---|---|---|---|---|---|---|---|---|---|---|---|---|---|
| | | | | | | Atopy* | EoE | | D | M | P | | |
| 1052 | 12 | M | W, BAA | OSHL | A, IgE-FA | Y | N | *unknown* | 35 | >40 | >40 | 4 | A |
| 478 | 21 | M | W | N | AR | N | N | milk, egg, legume | >40 | 0 | 5 | 5 | A |
| 489 | 14 | M | W | N | AD, AR | N | N | milk | 38 | 37 | VR | 3 | A |
| 8901 | 16 | F | W | N | *none* | Y | N | milk | >40 | 3 | 10 | NS | A |
| | | | | | | | | | 0 | 4 | 0 | 2 | R |
| 417 | 17 | M | W | N | AD, DA | Y | N | milk | >40 | 12 | 0 | 3 | A |
| 7528 | 15 | M | W | N | DA | N | N | milk | 10 | 12 | >40 | 3 | A |
| 9890 | 3 | M | W | N | AR, AD, DA, IgE-FA | Y | N | *unknown* | >40 | 35 | >40 | 3 | A |
| 474 | 11 | M | W | N | A, AR | Y | N | *unknown* | 4 | 20 | 0 | 3 | A |
| 7960 | 14 | M | W | N | AR | Y | N | *unknown* | >40 | 15 | 21 | 4 | A |
| 5276 | 3 | F | W | N | AD, IgE-FA | N | N | *unknown* | >40 | 3 | >40 | 3 | A |
| 5185 | 10 | M | W | N | *none* | Y | N | milk | 30 | 0 | 0 | 0 | A |
| 440 | 15 | M | W | N | A, AD, AR | Y | N | milk | 0 | VR | 0 | NS | R |
| 7687 | 8 | M | W | N | A | Y | N | milk | 0 | 0 | 0 | NS | R |
| 6398 | 9 | F | W | N | AR | Y | N | milk | 4 | 5 | 3 | 1 | R |
| 433 | 11 | M | W | N | DA | Y | N | milk | VR | VR | 0 | NS | R |
| 3295 | 16 | M | W | N | *none* | N | N | milk | 0 | 0 | 0 | 0 | R |

*eos/hpf, eosinophils per high-power field. D, distal. M, mid. P, proximal. EREFS, endoscopic reference score.*

*Atopy in family history defined as asthma, eczema, allergic rhinitis.

Key:

- Sex: M = male, F = female
- Race: W = white, BAA = Black/African American
- Ethnicity: N = not Hispanic/Latino, OSHL = other Spanish/Hispanic/Latino
- Allergic comorbidities: A = asthma, IgE-FA = IgE-mediated food allergy, AR = allergic rhinitis, AD = atopic dermatitis/eczema, DA = drug allergy, FPIES = food-protein-induced enterocolitis syndrome,
- Eosinophil count (eos/hpf): VR = “very rare”
- EREFS: scores reported as edema/rings/exudates/furrows/strictures = total, NS = not scored
- Group: A = active EoE, R = EoE in remission after diet therapy, N = not EoE

**Table S8. Fluorescently-labeled antibody panel used in esophagus flow cytometry data.**

| Marker | Fluorochrome | Clone | Manufacturer |
|---|---|---|---|
| Zombie Violet Live/Dead | | - | Biolegend |
| CD3 | PerCP-Cy5.5 | SP34-2 | BD Biosciences |
| CD4 | BV711 | RPTA-4 | Biolegend |
| CD45RA | BV605 | HI100 | Biolegend |
| CRTh2 | PE | HP-3G10 | Biolegend |
| CD161 | BV510 | BM16 | Biolegend |
| GPR15 | APC | SA302A10 | Biolegend |
| CD38 | BV650 | HB-7 | Biolegend |
| *CCR9*†* | *PE-Dazzle 594* | *LO53E8* | *Biolegend* |
| *CLA* | *PE-Cy7* | *HECA-452* | *Biolegend* |
| *α4β7* | *AlexaFluor488* | *Hu117* | *R&D Systems* |

*Analysis of markers in *italics* is not included in this report.

†For one sample in the Active group, the antibody cocktail did not include α4β7, CLA, CCR9.

**Table S9. Area under the curve (AUC) values for receiver-operator-characteristic curves.**

| Groups | Factor | AUC |
|---|---|---|
| Active vs Not EoE | GPR15 gMFI on GPR15+ peTh2 cells | 0.93 |
| | %GPR15+ peTh2 cells | 0.84 |
| | %CD38+ GPR15+ peTh2 cells | 0.78 |
| | GPR15 gMFI on GPR15+ memory CD4+ T cells | 0.71 |
| | %GPR15+ memory CD4+ T cells | 0.69 |
| | %CD38+ GPR15+α4β7+ peTh2 cells | 0.69 |
| Active vs Remission EoE | %CD38+ GPR15+CLA+ peTh2 cells | 0.74 |
| | %CD38+ GPR15+ peTh2 cells | 0.73 |
| | %CD38+ GPR15+α4β7+ peTh2 cells | 0.68 |

*EoE, eosinophilic esophagitis. gMFI, geometric mean fluorescence intensity. peTh2, pathogenic effector Th2.*

**Table S10. N's for main flow cytometry figures and reasons for sample exclusion.**

| Figure | Active | Remission | Not EoE | Samples excluded for |
|---|---|---|---|---|
| 1B | 29 | 26 | 16 (3 excluded) | CRTh2 staining difference (at room temperature in excluded samples) affecting # of positive cells |
| 1C | 29 | 26 | 19 | -- |
| 1D | 29 | 26 | 19 | -- |
| 1E, 4D, 6A | Memory CD4 – 29<br>Treg – 29<br>convTh2 – 28 (1 excluded)<br>peTh2 – 28 (1 excluded) | Memory CD4 – 26<br>Treg – 26<br>convTh2 – 26<br>peTh2 – 26 | Memory CD4 – 19<br>Treg – 16 (3 excluded)<br>convTh2 – 19<br>peTh2 – 17 (2 excluded) | - No FoxP3 staining (3 not EoE)<br>- Inadequate number of GPR15+ cells (<5) |
| 2A | 29 | 26 | 19 | -- |
| 2B, 6C | convTh2 – 28 (1 excluded)<br>peTh2 – 28 (1 excluded) | convTh2 – 26<br>peTh2 – 26 | convTh2 – 19<br>peTh2 – 17 (2 excluded) | Inadequate number of GPR15+ cells (<5) |
| 2C | 14 pairs (1 excluded) | | -- | Inadequate number of GPR15+ cells in active sample (<5) |
| InIN2E | a4b7+ – 29<br>a4b7+GPR15+ - 27 (2 excluded)<br>CCR9+ – 28 (1 excluded)<br>CCR9+GPR15+ - 14 (15 excluded)<br>CLA+ - 29<br>CLA+GPR15+ - 23 (6 excluded) | a4b7+ – 26<br>a4b7+GPR15+ - 24 (2 excluded)<br>CCR9+ – 24 (2 excluded)<br>CCR9+GPR15+ - 11 (15 excluded)<br>CLA+ – 26<br>CLA+GPR15+ - 19 (7 excluded) | a4b7+ – 19<br>a4b7+GPR15+ - 14 (5 excluded)<br>CCR9+ – 18 (1 excluded)<br>CCR9+GPR15+ - 6 (13 excluded)<br>CLA+ - 19<br>CLA+GPR15+ | Inadequate cell number positive for marker combination (<5) |
| 2F | convTh2s – 25 pairs (4 excluded)<br>peTh2s – 26 pairs (3 excluded) | convTh2s – 19 pairs (7 excluded)<br>peTh2s – 19 pairs (7 excluded) | convTh2s – 13 (6 excluded)<br>peTh2s - | Inadequate cell number (<5) in one or more pair members |
| 4C<br>(top, %AhR+) | *AhR+*<br>Memory CD4 – 29<br>Treg – 29 | *AhR+*<br>Memory CD4 – 26<br>Treg – 26 | *AhR+*<br>Memory CD4 – 16 (3 excluded)<br>Treg – 16 (3 excluded) | - No FoxP3 or AhR staining (3 not EoE)<br>- Inadequate number of cells in subset (<5) |

| | | | | |
|---|---|---|---|---|
| | convTh2 – 22 (7 excluded)<br>peTh2 – 25 (4 excluded)<br>*AhR-*<br>Memory CD4 – 29<br>Treg – 29<br>convTh2 – 28 (1 excluded)<br>peTh2 – 29 | convTh2 – 24 (2 excluded)<br>peTh2 – 19 (7 excluded)<br>*AhR-*<br>Memory CD4 – 26<br>Treg – 26<br>convTh2 – 26<br>peTh2 – 26 | convTh2 – 11 (8 excluded)<br>peTh2 – 10 (9 excluded)<br>*AhR-*<br>Memory CD4 – 16 (3 excluded)<br>Treg – 16 (3 excluded)<br>convTh2 – 16 (3 excluded)<br>peTh2 – 16 (3 excluded) | |
| 4C<br>(bottom, GPR15 MFI in AhR+ cells) | *GPR15+AhR+*<br>Memory CD4 – 28 (1 excluded)<br>Treg – 20 (9 excluded)<br>convTh2 – 9 (20 excluded)<br>peTh2 – 8 (21 excluded)<br>*GPR15+AhR-*<br>Memory CD4 – 29<br>Treg – 29<br>convTh2 – 28 (1 excluded)<br>peTh2 – 28 (1 excluded) | *GPR15+AhR+*<br>Memory CD4 – 26<br>Treg – 16 (10 excluded)<br>convTh2 – 6 (20 excluded)<br>peTh2 – 4 (22 excluded)<br>*GPR15+AhR-*<br>Memory CD4 – 26<br>Treg – 26<br>convTh2 – 26<br>peTh2 – 26 | *GPR15+AhR+*<br>Memory CD4 – 16 (3 excluded)<br>Treg – 6 (11 excluded)<br>convTh2 – 3 (16 excluded)<br>peTh2 – 1 (18 excluded)<br>*GPR15+AhR-*<br>Memory CD4 – 16 (3 excluded)<br>Treg – 16 (3 excluded)<br>convTh2 – 16 (3 excluded)<br>peTh2 – 14 (5 excluded) | - No FoxP3 or AhR staining (3 not EoE)<br>- Inadequate number of cells in subset (<5) |
| 5C | 11 | 6 | -- | -- |
| 5D | Memory CD4 – 11<br>convTh2 – 11<br>peTh2 – 11 | Memory CD4 – 6<br>convTh2 – 5 (1 excluded)<br>peTh2 – 4 (2 excluded) | -- | Inadequate number of GPR15+ cells (<5) |
| 5E | 10 pairs (1 excluded) | 1 pair (not shown, 5 excluded) | -- | Inadequate cell number (<5) in one or more pair members |

**Table S11. Fluorescently-labeled antibody panel used for sorting prior to single-cell RNA sequencing.**

| Marker | Fluorochrome | Clone | Manufacturer |
|---|---|---|---|
| Zombie Violet Live/Dead | | - | Biolegend |
| CD3 | PerCP-Cy5.5 | UCHT1 | BD Biosciences |
| CD4 | BV711 | SK-3 | BD Biosciences |
| CD45RA | BV605 | HI100 | BD Biosciences |
| CRTh2 | PE | HP-3G10 | BD Biosciences |
| CD161 | BV510 | BM16 | Biolegend |
| GPR15 | APC | SA302A10 | Biolegend |
| CD38 | BV650 | HB-7 | Biolegend |
| α4β7 | AlexaFluor488 | Hu117 | R&D Systems |
